# Evolution of degrees of carnivory and dietary specialization across Mammalia and their effects on speciation

**DOI:** 10.1101/2021.09.15.460515

**Authors:** Matthew D. Pollard, Emily E. Puckett

## Abstract

Conflation between omnivory and dietary generalism limits ecological and evolutionary analyses of diet, including estimating contributions to speciation and diversification. Additionally, categorizing species into qualitative dietary classes leads to information loss in these analyses. Here, we constructed two continuous variables – degree of carnivory (i.e., the position along the continuum from complete herbivory to complete carnivory) and degree of dietary specialization (i.e., the number and variety of food resources utilized) – to elucidate their histories across Mammalia, and to tease out their independent contributions to mammalian speciation. We observed that degree of carnivory significantly affected speciation rate across Mammalia, whereas dietary specialization did not. We further considered phylogenetic level in diet-dependent speciation and saw that degree of carnivory significantly affected speciation in ungulates, carnivorans, bats, eulipotyphlans, and marsupials, while the effect of dietary specialization was only significant in carnivorans. Across Mammalia, omnivores had the lowest speciation rates. Our analyses using two different categorical diet variables led to contrasting signals of diet-dependent diversification, and subsequently different conclusions regarding diet’s macroevolutionary role. We argue that treating variables such as diet as continuous instead of categorical reduces information loss and avoids the problem of contrasting macroevolutionary signals caused by differential discretization of biologically continuous traits.

---

Diet is a powerful selective force acting during evolution, with consequences for a species’ physiology, morphology, behavior, persistence time, and diversification rate. Examples of physiological traits shaped by diet include detoxification capacity (Robbins et al. 1991; Haywood et al. 2005), digestive enzyme production (Janiak et al. 2018), essential nutrient requirements (Knopf et al. 1978), and taste receptor function (Zhao et al. 2010; Jiang et al. 2012; Liu et al. 2016). Morphological adaptations that have arisen in response to diet vary in scale from small alterations that modify bite force (Santana et al. 2010; Law et al. 2018), to the extreme modifications observed in anteaters (Naples 1999) and mysticete whales (Deméré et al. 2008). High trophic level and narrow dietary breadth have been implicated in increased extinction risk (e.g., Purvis et al. 2000; Boyles and Storm 2007), leading to reduced species durations (Balisi et al. 2018).

Earlier work investigated the effect of diet on macroevolutionary dynamics. Price et al. (2012) estimated differential diversification rates between herbivores, omnivores, and carnivores. Specifically, omnivores had the lowest diversification rates while herbivores had the highest, and transitions into omnivory were more frequent than into either other category. Similarly, herbivorous birds (Burin et al. 2016), insects (Wiens et al. 2015), and crustaceans (Poore et al. 2017) also show elevated diversification rates. Birds show a similar pattern of low omnivore diversification rate, but a high rate of transitions into omnivory (Burin et al. 2016). Omnivory has thus been posited as a macroevolutionary sink, with the suggestion that this pattern may be widespread among higher taxonomic levels of tetrapods.

The effects of ecological factors on mammalian taxonomic diversification change across phylogenetic levels, producing clade- and level-specific diversification dynamics (Machac et al. 2017). At lower taxonomic levels, increasing specialization for frugivory was associated with higher diversification rates in bats (Rojas et al. 2012), but a diet-dependent diversification dynamic was not observed in murine rodents (Alhajeri and Steppan 2018). In ruminants, generalist herbivores showed higher diversification rates than browse-specialists (Cantalapiedra et al. 2014), which contradicts the prior notion of a positive relationship between dietary specialization and diversification rate in mammals. The conflicting results of the previous studies may be reconciled by the understanding that diet is capable of varying diversification relationships at different taxonomic levels.

Categorizing species into qualitative classes based on diet – especially using a coarse classification system of herbivore, carnivore, and omnivore – leads to a loss of information (Pineda-Munoz and Alroy 2014). Species with highly varied diets are clustered into potentially uninformative groupings. An additional problem with using a categorical variable is the need to choose diet categories and set boundaries between them, resulting in a lack of standardization of diet categorization in the literature. Between different studies, authors set conflicting cutoffs for clustering species. For example, one classification system separates mammals into hypercarnivores (>70% meat), mesocarnivores (50-70% meat), and hypocarnivores (>70% nonvertebrate foods) (Van Valkenburgh 1988, 2007). However, another system classified carnivores as those consuming only animal-based foods, herbivores as those consuming only plant-based foods, and omnivores as everything in between (Price et al. 2012). Other studies expand the number of different categories, further subdividing diet (e.g., Cantalapiedra et al. 2014; Burin et al. 2016). It requires great effort to construct a dietary dataset, as relevant data are dispersed across an extensive collection of primary literature or are unavailable for target species and must be generated. Depending on the research goals, studies may sacrifice detailed diet classification in exchange for broader taxonomic coverage, or vice versa. Comparisons between different studies are therefore made difficult by disagreement in how species should be grouped. A quantitative, continuous classification of diet based on food consumption percentages may be preferred, as downstream results are less impacted by arbitrary cutoffs decided upon earlier in the study (Pineda-Munoz and Alroy 2014).

Omnivory and dietary generalism are distinct yet conflated in some of the literature. Omnivores necessarily show more dietary generalism than an organism specialized to consume a single food type, yet this does not preclude herbivores or carnivores from consuming a broad range of foodstuffs. For example, a strictly herbivorous species (i.e., not an omnivore) may consume several distinct plant-based food items, making it a dietary generalist. Thus, we distinguish the macroevolutionary effects of degree of dietary specialization as independent of the effects of degree of carnivory (i.e., where a species is positioned on the continuum from complete herbivory to complete carnivory).

The independent macroevolutionary contributions of the degree of carnivory and degree of dietary specialization may vary in direction and extent across phylogenetic levels. In this study, our objectives are three-fold. First, we elucidate the history of diet across Mammalia by identifying shifts in the evolution of degrees of carnivory and dietary specialization. Second, we infer the relative importance of the degree of carnivory and degree of dietary specialization to evolution in different mammalian clades. Third, we identify and characterize significant relationships between diet and speciation rate across different taxonomic levels in Mammalia.

## Materials & Methods

### Preparation of Dietary Data and Tree Pruning

We utilized the EltonTraits dataset (Wilman et al. 2014) containing dietary data for 5,400 extant mammals. This data came primarily from Walker’s Mammals of the World (Nowak 1999), which provides accounts of species’ diets based on summaries of the existing literature. Qualitative descriptions of dietary preferences were translated by EltonTraits into numerical values representing the proportion of diet comprising specific food types. If a dietary description could not be obtained for a species, the proportion values were interpolated based on those from the same genus or family. In this study, the number of food types was condensed to six categories, giving the proportions of invertebrates, vertebrates, fruit, seeds, nectar, and other plant material (e.g., leaves).

We cross-referenced the species names in EltonTraits with those in the consensus version of the most recent mammal phylogeny (n = 4,098) (Upham et al. 2019). Mismatched and outdated species names were resolved using the *get_gbif_taxonomy* function of the traitdataform R package (Schneider 2018). We removed species not included in both datasets. We pruned the phylogeny using the *drop.tip* function of the ape R package (Paradis and Schliep 2018), leaving 3,649 species shared between the phylogeny and dietary dataset.

### Construction of “Degree of Carnivory” Variable

We built a continuous variable to represent the degree of carnivory. We first summed the proportions of fruit, seed, nectar, and other plant material into a single value representing the proportion of plant matter. Because the proportion variables sum to 100% and are therefore inappropriate for use in a principal component analysis (PCA), we transformed them into z-scores (Fraser et al. 2018). To convert the proportion of invertebrates, vertebrates, and plant matter into a continuous scale, we used discriminant analysis of principal components (DAPC) using the adegenet R package (Jombart and Ahmed 2011). The separation between groups is maximized during DAPC compared to (PCA) (Jombart et al. 2010). At two clusters, carnivores and herbivores were discriminated, while a third cluster distinguished extreme vertivores and invertivores (Fig. S1). In at least some clades, vertivory is considered an extreme form of carnivory, beyond that of invertivory (e.g., Giannini and Kalko 2005). Some researchers have considered vertivory as distinct from invertivory as it is from herbivory (Faurby et al. 2018). When invertivory and vertivory were separated using three clusters, vertivorous species obtained a marginally greater value for the first discriminant function than invertivorous species. This aligns with the idea that vertivory is a more extreme form of carnivory. Thus, we retained the DAPC results using three clusters. The resulting first discriminant function value was rescaled to a 0 – 100 scale, where 0 signifies complete herbivory and 100 complete carnivory.

### Construction of “Degree of Dietary Specialization” Variable

We made a second continuous variable to represent dietary specialization. We generated three new variables from the EltonTraits dataset that represent dietary breadth. First, we extracted the percentage (from 20 – 100%) of a species’ most common food type from the original six categories. Second, we summed the number of food types (from 1 – 6) included in each species’ diet. Third, to represent trophic breadth, we gave species that consumed only plants or animals a score of 1, and those that consumed both food types were given a score of 2. Although we argue that dietary generalism is distinct from omnivory, we propose that strictly herbivorous or carnivorous generalism allows for more specialized dietary adaptations than an omnivorous diet. Such adaptations can impact speciation rate and should be investigated. This informed our decision to incorporate trophic breadth as a component in our dietary specialization metric. We performed a PCA on these three variables in R using the MASS package (Venables and Ripley 2002), then rescaled the first principal component (PC1) to represent the degree of dietary specialization. A specialization score of 0 signifies a complete dietary generalist, whereas a 100 represents a complete dietary specialist.

### Regime Shift Analyses

Regime shifts represent changes in phenotypic optima on a phylogenetic tree under an Ornstein-Uhlenbeck (OU) model of trait evolution. We identified shifts during the evolution of carnivory and dietary specialization across Mammalia using the ℓ1ou method implemented in the l1ou R package (Khabbazian et al. 2016). The ℓ1ou method fits a multi-optimum OU model of trait evolution using a phylogenetic lasso approach and does not require prior hypotheses regarding the location of shifts. OU models assume random change in a trait value across the tree, but with a centralizing tendency back to an optimum value (Butler and King 2004). A multi-optimum OU model assumes the presence of multiple regime shifts toward different trait optima throughout the phylogeny. The ℓ1ou method detects the scale and location of shifts in the optimum trait value.

We used a Bayesian Information Criterion (BIC) approach to select the best-fitting configuration of evolutionary regimes. To reduce the number of low probability shifts detected while using the BIC, we reduced the maximum number of shifts allowed to *n*/10, where *n* was the number of tips in the tree. This represents a compromise between the default maximum of *n*/2 and the conservative maximum of √*n* + 5 recommended by alternative regime shift analysis methods (Bastide et al. 2018).

The ℓ1ou method can handle large phylogenetic trees comprising thousands of taxa (Khabbazian et al. 2016); however, the analyses did not complete when using our pruned mammal phylogeny as input (analysis cancelled after 28 d with 24 CPUs). Therefore, we reduced the number of species by first removing those with diets inferred via interpolation in the original EltonTraits dataset. Then we subset the tree so that, for each genus, a maximum of three species could share the same carnivory or dietary specialization score. This produced a tree of 1,912 species for our analysis of regime shifts in degree of carnivory, and a tree of 1,941 species for degree of dietary specialization. This maximized representation of dietary diversity in each genus while still producing input trees of a suitable size for ℓ1ou.

### Phylogenetic Half-lives of Degrees of Carnivory and Dietary Specialization

The phylogenetic half-life (t_1/2_) of a trait represents the time it takes for a species entering a new niche or regime to evolve halfway to the new optimum trait value (Hansen 1997). The α parameter returned by each ℓ1ou analysis represents selection strength and was used to calculate phylogenetic half-life for degrees of carnivory and dietary specialization. Because ℓ1ou rescales the phylogenetic tree height to one, the calculated t_1/2_ values are percentages of the original clade age (Ho and Ané 2014). A short t_1/2_ relative to tree height indicates that evolution towards new optimum carnivory or dietary specialization scores was fast, with weak residual phylogenetic correlations. Conversely, if t_1/2_ was large relative to tree height, the strength of the OU process was weak and may be indistinguishable from Brownian motion (Cooper et al. 2016). Indeed, an α value of zero indicates that the trait did not evolve under an OU process in that clade (Hansen et al. 2008; Uyeda and Harmon 2014).

### Continuous Trait-dependent Speciation Rates

We characterized the impact of degree of carnivory and dietary specialization on speciation using the quantitative state speciation and extinction (QuaSSE) method in the diversitree R package (FitzJohn 2010, 2012). For both traits, we constructed various models, each fitting a different relationship between the trait of interest and speciation rate, including: a constant speciation rate independent of the trait value, a linear relationship between trait value and speciation rate, a sigmoidal relationship, and a unimodal relationship. Given known difficulties with extinction rate detection in QuaSSE (FitzJohn 2010) and broader issues with the estimation of extinction from molecular phylogenies (Rabosky 2010), we decided to fit a constant extinction rate instead of allowing the rate to vary with the trait (as in Velasco et al. 2016). Thus, if we observed an effect of diet on diversification, it was due to a diet-dependent speciation dynamic. We selected the best-fitting model using the Akaike information criterion (AIC). To account for the influence of phylogenetic uncertainty on our results (Pagel and Lutzoni 2002), we first performed these analyses using the pruned consensus phylogeny, then repeated the analyses on 10 trees sampled from a credible set of 10,000 DNA-only mammalian phylogenies produced by Upham et al. (2019). These trees were pruned in the same way as the consensus tree (see above). To understand how phylogenetic level affects the relationship between diet and speciation rate, we fit QuaSSE models for Mammalia and nine lower clades: Artiodactyla + Perissodactyla, Atlantogenata, Carnivora + Pholidota, Chiroptera, Euarchonta, Eulipotyphla, Lagomorpha, Marsupialia, and Rodentia.

We evaluated each QuaSSE model with a simulation approach (as in Inostroza-Michael et al. 2018) to address concerns of high Type 1 error rates (i.e., that neutral traits are significantly associated with diversification rate (Rabosky and Goldberg 2015)). We used the *fastBM* function in the phytools R package (Revell 2012) to simulate random traits on each phylogeny 100 times under a model of Brownian motion. For these simulations, we used the empirically estimated diffusion parameters from the QuaSSE models that fitted a constant relationship between speciation rate and trait. The ancestral root values for the simulations were estimated using the *fastAnc* function in phytools. We fit the constant model and the best-fitting model from the observed data to each set of simulated trait values and calculated log-likelihood differences between the pairs of models. A *P*-value was created using the proportion of simulated values that achieved a log-likelihood difference as large as that obtained in the observed data. Because we used multiple phylogenies in our QuaSSE analyses to address phylogenetic uncertainty, we considered a clade’s diet-dependent speciation dynamic to be very robust to uncertainty if the *P*-value of the best fitting QuaSSE model was ≤ 0.05 in at least ten out of the eleven phylogenies tested. We considered the dynamic only moderately robust if *P* ≤ 0.05 in eight out of eleven phylogenies, and not robust to phylogenetic uncertainty if *P* ≤ 0.05 for less than eight trees. To visualize the overall relationship between trait and rate for each speciation dynamic, we averaged across the best-fitting models.

### Categorical Trait-dependent Diversification Rates

Our work is not directly comparable to previous studies since we created a synthetic continuous variable instead of using categories. Thus, to compare our results, we used the multiple state speciation and extinction (MuSSE) method in the diversitree R package (Fitzjohn 2012) to characterize diet-dependent diversification using different qualitative categorizations of diet. We categorized diet in two ways; first, we used the species and groupings in Price et al. (2012) (we will refer to this as Price categorization). There, data was based on literature searches of direct dietary composition from gut, feces, or observational data, which was then binned into plants only (herbivore), plant and animal (omnivore), and animal matter only (carnivore). Second, we replicated the Price categorization but used the raw EltonTraits data (and not our continuous carnivory score), where species with diets composed of 100% animal matter were carnivores, 0% animal matter were herbivores, and 1 – 99% animal matter were omnivores (we will refer to this as Replicated categorization). We limited our MuSSE analyses to species included in both the Price categorization and EltonTraits data (n = 1,323).

## Results

### Constructed Dietary Variables

We created a synthetic variable to quantify degree of carnivory ranging from completely herbivorous (0) to completely carnivorous (100; Fig. 1). Notably, herbivorous dietary guilds (folivores, frugivores, granivores, and nectarivores) had a median score of 0. The median carnivory scores for invertivores and vertivores were 87 and 95, respectively, with overlap between the two dietary guilds. Although both groups are considered carnivorous, the constructed variable attributed a higher degree of carnivory to a vertebrate-consuming species than a species whose diet comprises an equivalent proportion of invertebrates. Species with carnivory scores of 100 were therefore completely vertivorous, an artifact from the variable construction (Fig. S1).

**Figure 1.**
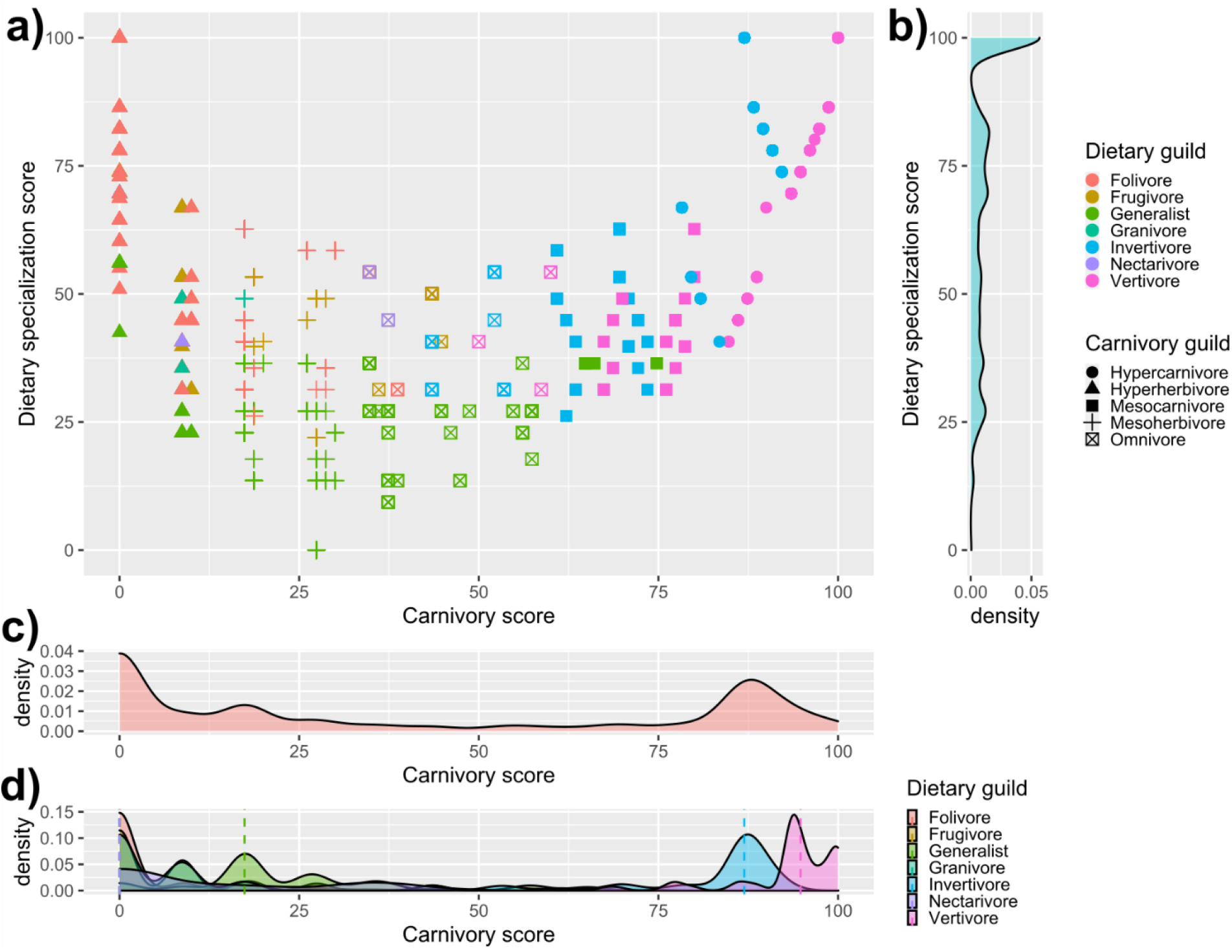
a) The relationship between constructed variables measuring the degree of carnivory and degree of dietary specialization. To visualize how species (n = 3,649) were distributed across the variable space, they were grouped into dietary guilds. Species were assigned to a specific dietary guild if their diet comprised ≥50% of that food type. If no food type comprised ≥50% of a species’ diet, they were assigned to the generalist guild. Hypercarnivores are categorized as species that consume ≥80% animal matter in their diet, mesocarnivores consume 60 – 80% animal matter, omnivores consume 40 – 60% animal matter, mesoherbivores consume 20 – 40% animal matter, and hyperherbivores consume ≤20% animal matter. b) Distribution of species along the dietary specialization spectrum. c) Distribution of species along the carnivory spectrum. d) Distribution of dietary guilds along the carnivory score spectrum.

We constructed a variable representing the degree of dietary specialization using PC1, which explained 83.2% of the variance (Fig. S2). Lower dietary specialization scores indicated increased dietary generalism. Species with scores of 100 had diets composed of a single food type, whereas 0 represented a diet comprising six food types across trophic levels, with the most abundant category comprising 20% of the diet. Dietary specialization showed a convex curvilinear relationship with carnivory, indicating that highly specialized species were also extreme herbivores or extreme carnivores. Because of the higher number of herbivore dietary guilds in EltonTraits, the relationship between the degrees of carnivory and dietary specialization was skewed to the right (Fig. 1a).

### Regime Shifts in Degrees of Carnivory and Dietary Specialization across Mammalia

We detected 134 and 50 regime shifts in degree of carnivory and dietary specialization across Mammalia, respectively (Fig. 2–3; Table S1). In the analysis of carnivory score, the regime occupied by the root had an optimal score indicative of a diet comprising 80-90% animal matter, similar to extant species with a substantial invertebrate component. This original regime is still occupied by extant members of Carnivora, Atlantogenata, and Marsupialia. In the analysis of dietary specialization, the root occupied a regime with a highly specialized optimum diet. This regime is still occupied by members of most mammalian clades, excluding Marsupialia and Rodentia. The phylogenetic half-life of degree of carnivory was 27% of the half-life of degree of dietary specialization (Table S2), indicating that carnivory score shows a lower phylogenetic signal. Thus, compared to dietary specialization scores, there is less statistical dependence among species’ carnivory scores due to phylogenetic relatedness (Revell et al. 2008). Regime shifts varied in frequency across the phylogeny. We provide detailed, clade-specific descriptions and interpretations of regime shift results in the Supporting Information (Figs. S11-S19).

**Figure 2.**
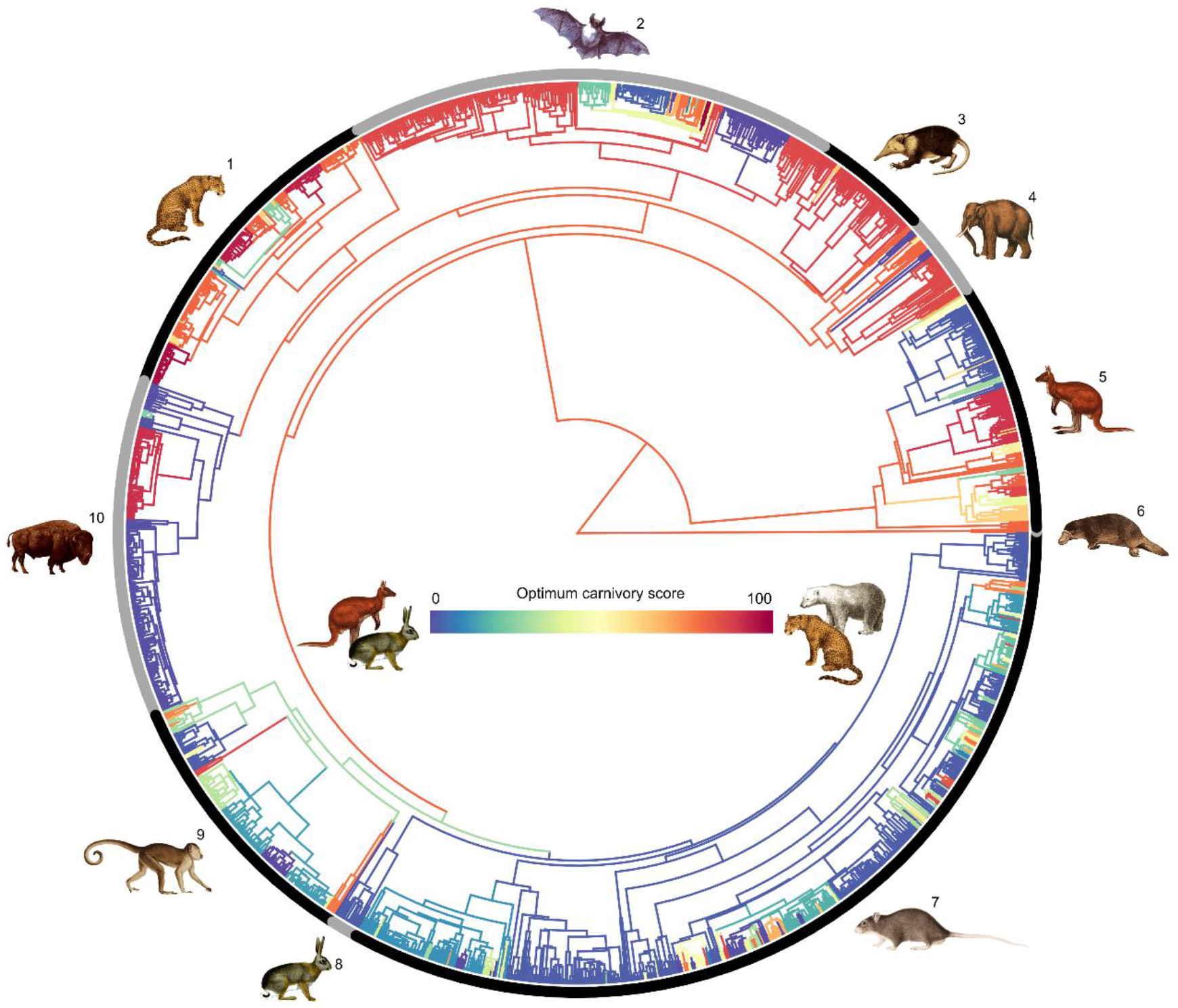
Regime shifts in optimum carnivory score in Mammalia. Branch colors indicate the value of the optimum carnivory score, with blue indicating a low score, and red indicating a high score. The positions of branch color changes show the location of the regime shifts. A gray branch represents lack of data regarding regime occupancy for that branch because it was not included in any regime shift analyses. The numbered black and gray bars encircling the phylogeny indicate the individual clades on which QuaSSE analyses were performed. 1, Carnivora + Pholidota; 2, Chiroptera; 3, Eulipotyphla; 4, Atlantogenata; 5, Marsupialia; 6, Monotremata; 7, Rodentia; 8, Lagomorpha; 9, Euarchonta; 10, Artiodactyla + Perissodactyla. Images source: Wikimedia Commons.

**Figure 3.**
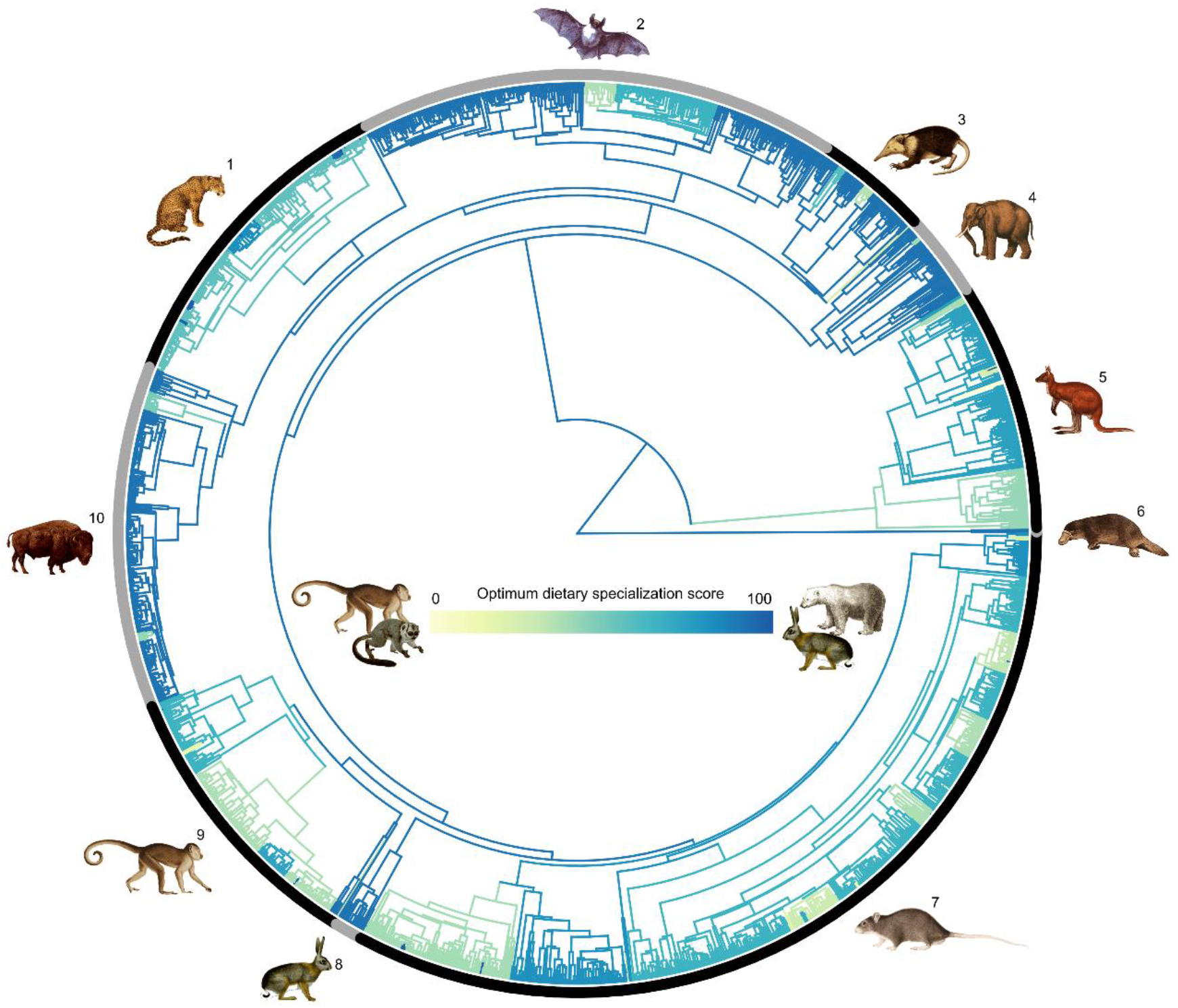
Regime shifts in dietary specialization score in Mammalia. Branch colors indicate the value of the optimum specialization score, with yellow indicating a low score, and blue indicating a high score. The positions of branch color changes show the location of the regime shifts. A gray branch represents lack of data regarding regime occupancy for that branch because it was not included in any regime shift analyses. The numbered black and gray bars encircling the phylogeny indicate the individual clades on which QuaSSE analyses were performed. 1, Carnivora + Pholidota; 2, Chiroptera; 3, Eulipotyphla; 4, Atlantogenata; 5, Marsupialia; 6, Monotremata; 7, Rodentia; 8, Lagomorpha; 9, Euarchonta; 10, Artiodactyla + Perissodactyla. Images source: Wikimedia Commons.

### Diet-dependent Speciation Rates

We tested if carnivory or dietary specialization was related to speciation rate within each clade using QuaSSE (Table 1). We found relationships between speciation rate and degree of carnivory that were very robust to phylogenetic uncertainty in Artiodactyla + Perissodactyla, Carnivora + Pholidota, Marsupialia, and Mammalia as a whole. Specifically, most significant QuaSSE models for Mammalia fitted an inverse unimodal relationship between trait and rate (Figs. 4 and S3). Across these models, speciation rate was lowest for intermediate carnivory scores and increased towards both extremes of the carnivory spectrum. However, speciation rates were higher for extreme herbivores than extreme carnivores. Artiodactyla + Perissodactyla, Carnivora + Pholidota, and Marsupialia also had inverse unimodal relationships (Figs. S4-S6). Two other clades, Chiroptera and Eulipotyphla, showed carnivory-dependent speciation dynamics that were only moderately robust to phylogenetic uncertainty. Chiroptera showed a similar relationship as previously described, while Eulipotyphla had a sigmoidal relationship where speciation rate showed a sharp increase at the hypercarnivorous end of the carnivory spectrum (Figs. S7-S8). The phylogenetic robustness of the other clades’ dynamics were low enough that we inferred there is no significant relationship between degree of carnivory and speciation rate in those clades. For dietary specialization, a robust association with speciation rate existed only for Carnivora + Pholidota (Fig. S9). Carnivora + Pholidota had a positive linear relationship where speciation rate was highest for complete dietary specialization and lowest for extreme dietary generalism, reaching zero lineages per Myr for specialization scores below 20. The best-fitting QuaSSE models for Atlantogenata and Lagomorpha were consistently the null, indicating that there was no relationship between speciation rate and either diet score in these clades.

**Table 1.** Diet-dependent speciation dynamics as inferred by simulation-based evaluation of best-fitting QuaSSE models using eleven mammal phylogenies. The number of statistically significant QuaSSE models (out of 11) is shown for each taxa and our two variables. Clades for which the best-fitting QuaSSE model was the null were not evaluated via simulations and are labelled as ‘NA’. *P*: the proportion of Brownian motion simulations that produced log-likelihood differences as large as obtained in the observed data.

| Clade | Carnivory | Dietary specialization |
| --- | --- | --- |
| | No. trees/models with $P \leq 0.05$ | No. trees/models with $P \leq 0.05$ |
| Mammalia | 10 | 3 |
| Artiodactyla + Perissodactyla | 11 | 0 |
| Atlantogenata | NA | NA |
| Carnivora + Pholidota | 11 | 11 |
| Chiroptera | 8 | 5 |
| Euarchonta | 1 | 0 |
| Eulipotyphla | 9 | 0 |
| Lagomorpha | NA | NA |
| Marsupialia | 11 | 6 |
| Rodentia | 6 | 3 |

**Figure 4.**
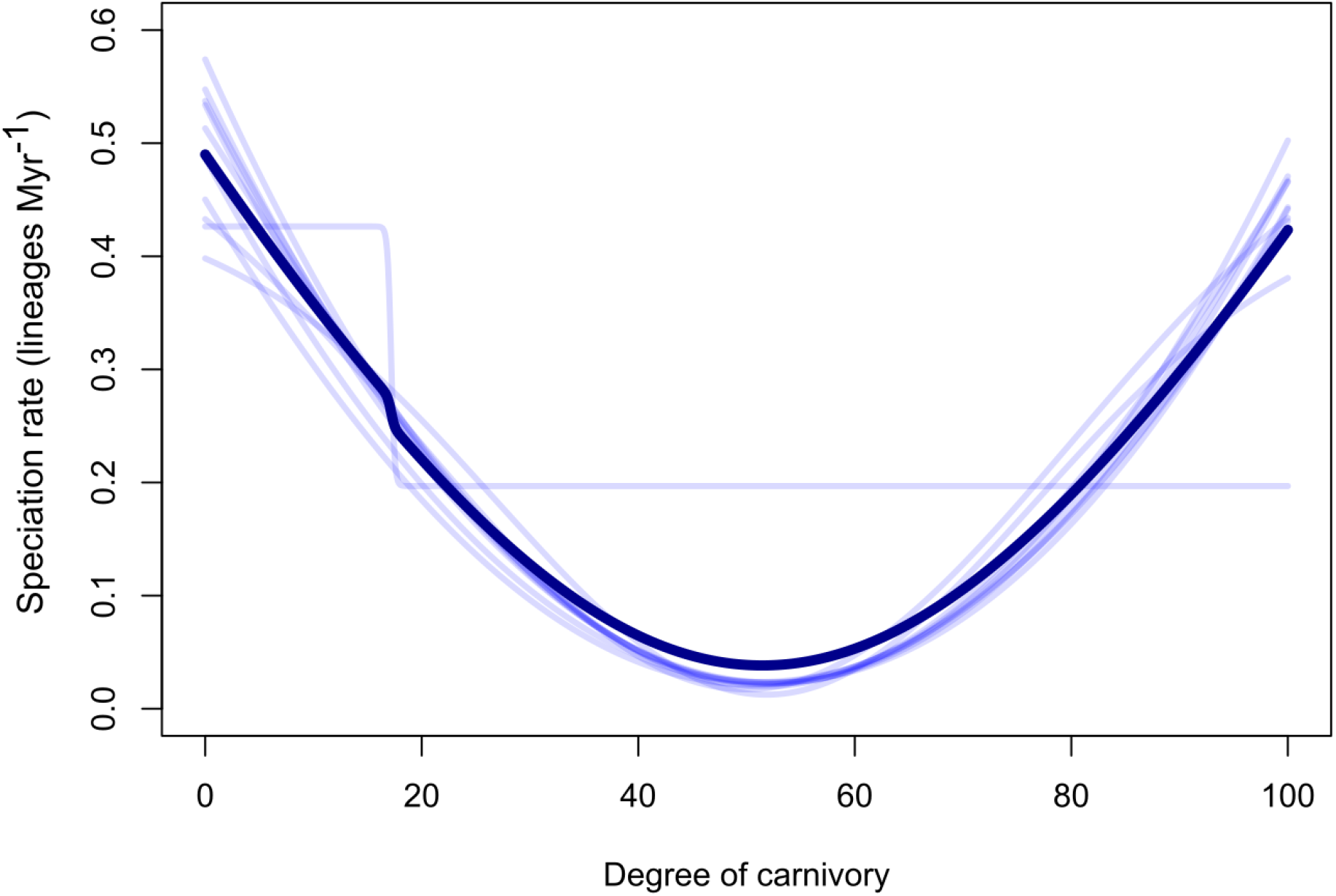
The relationship between degree of carnivory and speciation rate in Mammalia. Across the eleven trees used in this study, ten were associated with a statistically significant best-fitting QuaSSE model. The transparent lines show the relationships predicted by these models. The dark line represents the relationship produced by averaging over each significant model.

### Diet-dependent Diversification using Categorical Variables

Given our creation of a synthetic carnivory score, we decided to compare the results using our continuous scale to a previous categorical analysis. We observed that the diversification rate varied depending on how diet was classified (Fig. 5). Using the Price categorization, herbivores had the highest diversification rates, while omnivores had the lowest (Fig. 5a), concordant with previously published results. While we were able to reproduce the pattern from the previous study using the more recent phylogeny, our estimates of diversification rate were lower for each category compared to Price et al. (2012). However, when using the EltonTraits categorization, carnivores had the lowest diversification rates, while omnivores and herbivores had overlapping rates (Fig. 5b).

**Figure 5.**
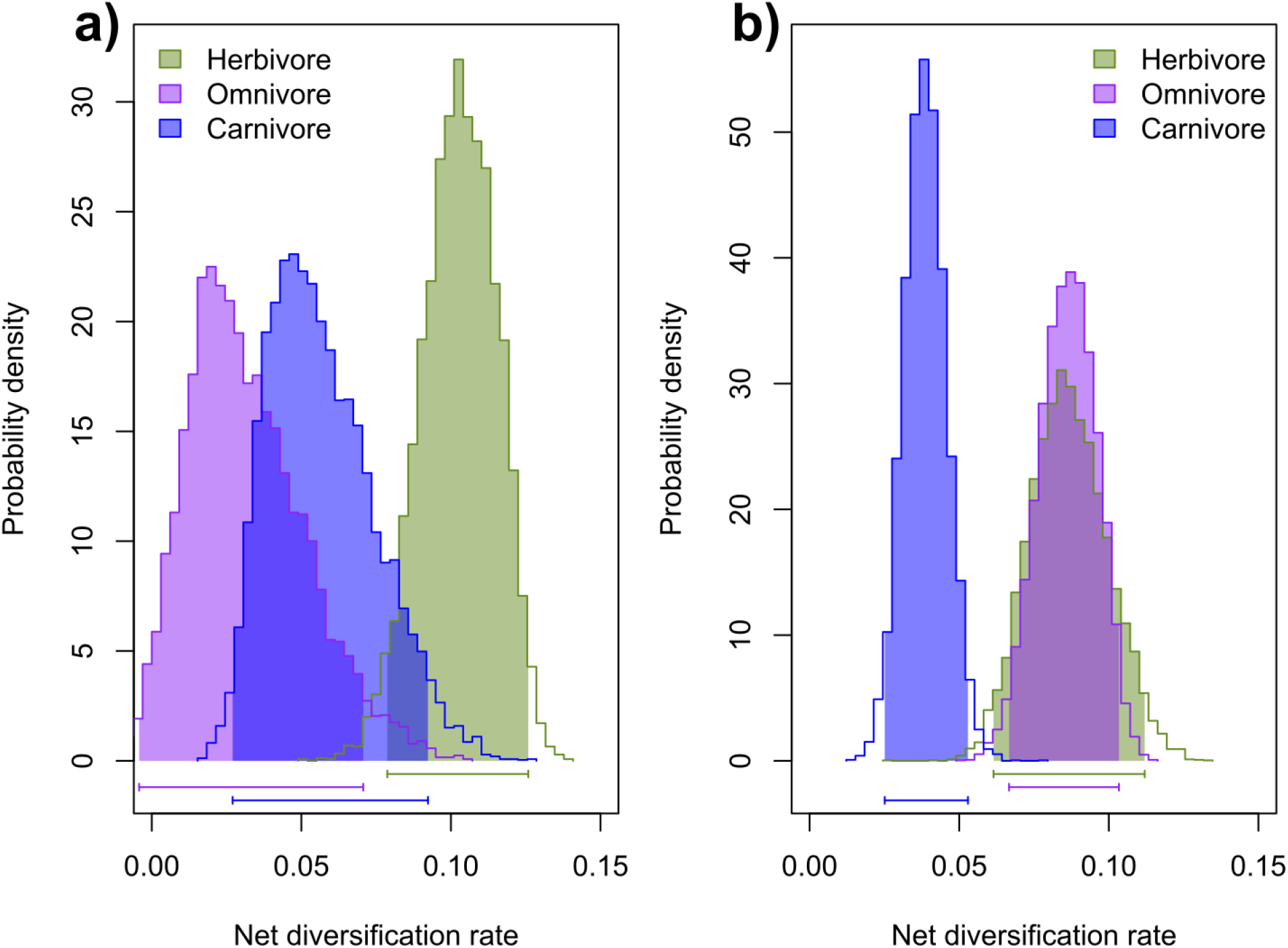
Posterior probability densities for net diversification rate of herbivores, omnivores, and carnivores (n = 1,323), using two diet classification schemes: a) The diet groupings of Price et al. (2012) were reused with the most recent mammalian phylogeny (Upham et al. 2019), b) The classification method of Price et al. (2012) replicated using EltonTraits data (animal matter: 0% herbivore, 1-99% omnivore, 100% carnivore).

Transition rates between dietary guilds also changed depending on classification system. Only the MuSSE analysis using the Price categorization reported highest transition rate from herbivory to omnivory (Fig. S10a). Analysis using our EltonTraits categorization reported highest transition rates from omnivory to herbivory (Fig. S10b).

## Discussion

### Mammalian Diversification is Shaped More by Degree of Carnivory than Dietary Specialization

Our analyses teased apart the independent contributions to mammalian diversification of degree of carnivory and degree of dietary specialization, two aspects of diet that have typically been conflated. We observed that the effect of diet on speciation varied with phylogenetic level, and that the selective importance of degrees of carnivory and dietary specialization differed among taxonomic groups (Table S2). Notably, degree of carnivory showed a statistically significant relationship with speciation rate when assessed across all of Mammalia (Fig. 4), as well as in at least three lower clades (Figs. S4-S8). Across mammals, omnivorous species had the lowest rates (Figs. 4 and S3), which is a pattern previously observed in mammals and birds (Price et al. 2012; Burin et al. 2016). Our findings show that this pattern can extend to clades at lower taxonomic levels (i.e., Artiodactyla + Perissodactyla, Carnivora + Pholidota, Marsupialia, Chiroptera, and Eulipotyphla), but that it does not describe all lower groupings. Conversely, there was no significant relationship between dietary specialization and speciation across Mammalia as a whole, and only Carnivora + Pholidota showed a significant impact of dietary specialization at lower taxonomic levels (Fig. S9). Thus, we find that diet does not have a homogeneous effect on diversification at different hierarchical levels of Mammalia. This is consistent with the level-dependent effects of other ecological factors that govern mammalian diversification (Machac et al. 2017).

Five of the clades we analyzed showed no significant diversification relationship with diet. Monotremata comprised only three species in our study, so it was not included in our analyses. Diets within Lagomorpha are relatively homogeneous, with few regime shifts in degree of carnivory or dietary specialization throughout the clade (Fig. S16). Thus, lagomorph speciation was unlikely to be shaped by variation in diet. Conversely, diets in Atlantogenata, Euarchonta, and Rodentia span the breadth of the carnivory spectrum, but these clades still showed no diet-dependent speciation dynamics. Lack of a relationship between a particular trait and speciation rate does not necessarily indicate that the trait is irrelevant to speciation, and by extension, diversification. While a trait may not be the rate-limiting control, it may still play an important role in the evolution of a clade (Rabosky 2015). Other aspects of an organism’s biology may be more important in shaping speciation and diversification. For example, locomotory mode was found to impact diversification of murine rodents, with less specialized modes associated with higher diversification rates (Nations et al. 2020).

Previous explanations for decreased diversification of omnivores have focused on the effect of dietary specialization on diversification rate, rather than the effect of degree of carnivory. Specialists show increased competitive ability over generalists when in their optimal environment (Wilson and Yoshimura 1994; Büchi and Vuilleumier 2014). Generalists utilize several resources instead of being most effective at using a single one, so will be excluded from the optimal environments of specialist species (MacArthur and Levins 1964). Such outperformance by specialists should have macroevolutionary costs for omnivores, resulting in low speciation and/or high extinction rates (Burin et al. 2016). Specialized species are more prone to allopatric speciation because their narrow niches cause fragmented distributions and isolate formation as environments change (Futuyma and Moreno 1988; Allmon 1992). Specialists also suffer resource limitations more frequently than generalists, so experience stronger directional selection in a changing environment (Vrba 1980, 1987). These explanations may apply to Carnivora + Pholidota, for which dietary generalism was associated with lower rates, and specialization with higher rates. However, in most clades and in Mammalia as a whole, dietary specialization did not have a significant relationship with diversification. As dietary generalism is not directly associated with lower rates in most clades, these suggested explanations are insufficient.

Reduced omnivore diversification is not due to a purely generalism-dependent speciation dynamic. Instead, consumption of both plant and animal matter causes lowered speciation rates in mammals. Future explanations for the role of omnivory in reducing diversification rate should seek to answer how differences in degree of carnivory lead to different speciation rates. Increased herbivore diversification may be explained by primary producers comprising the largest proportion of biomass on Earth (Bar-On et al. 2018), suggesting that more herbivorous species can be supported globally. Ecological factors limit diversity and species richness within a clade, and net diversification rate slows down over time in a diversity-dependent manner as this limit is reached (Rabosky 2009). By extension, herbivores should experience a greater net diversification rate than omnivores as it takes them longer to reach the diversity limit imposed on them. However, this explanation for increased herbivore speciation is plausible only if ecological limits to species richness occur. There is an unresolved debate over the existence of diversity-dependent regulation of species richness (Rabosky and Hurlbert 2015), with some arguing that such an equilibrial process does not exist, and that species diversity is unbounded at large scales (Harmon and Harrison 2015).

Extreme carnivory was also associated with high speciation rates, despite typically being associated with increased extinction (Purvis et al. 2000; Estes et al. 2011). This is unsurprising given that speciation and extinction rates are often correlated, with traits that promote speciation also increasing extinction risk (Stanley 1990; Greenberg and Mooers 2017). High rates in carnivores may be attributed to increased diversification of prey species at lower trophic levels, which drives evolution and diversification of predatory species in an upwards adaptive radiation cascade (Brodersen et al. 2018; Pontarp 2020).

The relationship between diet and speciation could be indirect. An unmeasured trait co-distributed with diet may be more responsible for shaping mammalian diversification (Beaulieu and O’Meara 2016). Our QuaSSE analyses cannot account for a hidden factor that affects speciation rate when the degree of carnivory or dietary specialization varies. For example, body size is associated with degree of carnivory in mammals, with herbivores having the largest masses and carnivores the smallest (Price and Hopkins 2015). Diversification rates are also higher for smaller species (Martin 2017); thus, body size may be an important factor related to increased carnivore speciation, but may not explain increased speciation across herbivores. Dispersal ability is also correlated with degree of carnivory, with carnivores having greater dispersal distances than omnivores and herbivores (Sutherland et al. 2000; Santini et al. 2013). Significant associations between dispersal ability and speciation rates have been observed in both mammals and birds (Claramunt et al. 2012; Upham et al. 2020), but the correlation between trait and rate is negative, with fastest rates in low-vagility species. Thus, the co-distribution of diet and dispersal ability could explain the high herbivore speciation rates observed in our study, but not those of carnivores.

If the link between degree of carnivory and speciation is indeed indirect, the patterns we observed may manifest through diet’s relationship with multiple traits, potentially with different mechanisms operating in different regions of the phylogeny. For example, in Carnivora and other mammalian lineages, there is an evolutionary trend of increasing body size over time, known as Cope’s Rule (Alroy 1998). Within Carnivora, there is an association between increasing body size and specialization for hypercarnivory, and this has been posited as a reason for bias towards evolution of hypercarnivory in the carnivoran fossil record (Van Valkenburgh et al. 2004; Van Valkenburgh 2007). Thus, Cope’s Rule may contribute to the positive relationship between increasing degrees of carnivory and speciation rate in Carnivora. In Marsupialia, environmental changes may contribute to the observed link between diet and speciation rate. Carnivorous dasyurids and herbivorous diprotodontids comprise much of the marsupial diversity. The radiation of these clades has been attributed to the aridification of Australia (Dodt et al. 2017; García-Navas et al. 2018), with later waves of diversification associated with subsequent major expansion of grasslands (Kealy and Beck 2017; Celik et al. 2019). Carnivory and herbivory in dasyurids and diprotodontids, respectively, may be a small part of a larger suite of adaptations to fill the new niche space created by aridification. Thus, the diet-dependent speciation dynamic observed in Marsupialia may have been reinforced or driven by the impact of environmental change on specific clades.

### The Optimum Diet at the Origin of Mammalia was Insectivorous

Through our evolutionary shift analyses, we inferred that the optimum dietary condition at the origin of Mammalia was biased towards insectivory. This result supports the consensus that the ancestral mammal was an insectivore (Kemp 2005). Extant species that occupy both the ancestral carnivory regime and the ancestral dietary specialization regime are restricted to Monotremata and within Xenarthra, in Cingulata and Vermilingua. The diets observed in these clades might provide a good estimate for the dietary composition of early mammals, but continued occupancy of the ancestral regimes through to present day is not an indication that early mammalian diets were identical. In OU models, shifts in phenotypic optima represent changes in the selective pressure acting on the trait of interest. These shifts can coincide with historical events such as habitat change or changes in other traits. Thus, a dietary shift at the origin of a clade offers insight into the events that formed the lineage. We provide an account of regime shifts in degrees of carnivory and dietary specialization across Mammalia in our Supporting Information. This may serve as a valuable resource for researchers investigating the origin and evolution of specific mammalian clades.

### Limitations

Concerns over the reliability of our underlying data raise questions regarding the accuracy of our results. The EltonTraits dataset, used to construct our continuous dietary variables, was built primarily from qualitative descriptions in secondary sources instead of direct estimations of diet composition from gut or fecal sampling. These descriptions were translated into numerical values representing the proportion of diet comprising specific food types, then further rounded to the nearest decile. Additionally, for over 1,000 species in the dataset, the proportions were interpolated based on values from the same genus or family. A dataset compiled from quantitative data for each of the >5,600 mammalian species would both be a valuable community resource and strengthen the inferences of the current study. However, while it is possible to construct large datasets using such dietary data (e.g., Grundler 2020; Grundler and Rabosky 2021), these datasets require substantial time and effort to produce using standardized estimation approaches and are still limited to several hundred species, rather than the thousands that represent all of Mammalia. To analyze diet across almost all mammals, as in this study, we are currently limited to using comprehensive dietary datasets constructed using qualitative, verbal diet descriptions from secondary sources. Such datasets include EltonTraits and similar alternatives (Jones et al. 2009; Kissling et al. 2014). EltonTraits has helped answer impactful questions in ecology and evolution, including how diet affects the evolution of mammalian gut microbiomes (Groussin et al. 2017), whether diet affects species’ responses to climate change (Buckley et al. 2018), and how ecological and functional diversity is declining (Cooke et al. 2019; Brodie et al. 2021). Quantitative descriptions may be unavailable for many biologically continuous traits, but this should not prevent attempts to capture their continuous nature in future studies. As Price et al. (2012) showed before us, it is still possible to make important inferences about the role of diet on speciation and diversification while using qualitative underlying data.

The specialization score does not consider the full scope of dietary specialization because it is limited by the data used to construct it. The EltonTraits dataset differentiates between vertebrates, invertebrates, fruits, nectar, seeds, and other plant material in each species’ diet (Wilman et al. 2014), allowing us to construct our coarse estimation of dietary specialization. However, some species display extreme dietary specializations that are inadequately captured by broad food categories. For example, the giant panda is a bamboo specialist, feeding on specific parts of certain bamboo species (Wang et al. 2017). Despite this extreme taxonomic specialization, the panda’s diet in EltonTraits is coded similarly to species that feed on multiple, varied foodstuffs, such as grasses, lichen, shrubs, and fungi. This is because these foodstuffs all fall into the ‘other plant material’ category. We may expect more nuanced results for diet-dependent speciation rates if detailed dietary data was used to construct a synthetic specialization score. Thus, while this is the first effort to disentangle specialization from the carnivory-to-herbivory axis, significant refinements to the specialization score could be made, particularly for smaller phylogenies where complete dietary data may be accessible. We further acknowledge that intraspecific variation in niche specialization exists, and degree of individual specialization varies between species (Bolnick et al. 2003). However, each species represents a single entry in the EltonTraits dataset, meaning that population specialization cannot be considered. It is possible that intraspecific differences impact mammalian speciation, but this effect is impossible to detect in our analyses. Nevertheless, our findings indicate that dietary specialization measured at a broader scale does not impact speciation in most mammalian clades.

A recent study demonstrated that birth-death models cannot reliably estimate speciation and extinction rates on extant timetrees (Louca and Pennell 2020). For any diversification scenario, there are an infinite number of equally likely alternative scenarios. Despite potentially having very different diversification dynamics, these congruent scenarios cannot be distinguished from one another. Because state-dependent speciation and extinction (SSE) models such as QuaSSE are extensions of the birth-death model, it is possible that they do not accurately estimate the relationship between rates and traits. It is important to acknowledge this unidentifiability issue when considering the findings of our study, but also to note that it does not mean our results are inaccurate. We compared a small number of alternative models representing the potential relationship between diet and speciation rate for each clade. This was due necessarily to there only being four possible relationships that QuaSSE is capable of fitting (constant, linear, sigmoidal, and unimodal/humped). By only comparing these, we avoided the problem of there being many congruent models (Morlon et al. 2020). Instead, we infer that one of the models from our limited selection fits the data best and is the more plausible scenario of speciation. Additionally, by considering character state evolution, SSE models consider more than just the timing of branching events in lineage-through-time plots as in the models of Louca and Pennell (2020). This may allow SSE models to overcome the unidentifiability issue (Helmstetter et al. 2021). In any case, we were less concerned with estimating precise rate values and more interested in characterizing differences in speciation rate as diet changes. By analyzing multiple alternative trees to account for phylogenetic uncertainty and then averaging over the best-fitting models for each tree, we also account for uncertainty in model fit. In doing so, we strengthen our confidence in the accuracy of the diet-dependent speciation dynamics inferred by our analyses.

### Continuous vs Categorical Representations of Diet

Our analysis representing diet as a continuous variable was concordant with a previous study where diet was considered a categorical variable (Price et al. 2012); specifically, both studies observed that diversification was lowest for omnivores (Fig. 4). However, while converting the EltonTraits data into categorical carnivory bins for validation, we observed that diet-dependent diversification rates varied due to differences in dietary classification (Fig. 5). We were unable to reproduce results from Price et al. (2012) using our own categorical variable. Our analysis using the Replicated categorization also found that the highest transition rates between trophic strategies were out of omnivory and into herbivory or carnivory (Fig. S10b). This is the opposite result to Price et al. (2012) and to our analysis using the Price categorization, where transitions into omnivory had the highest rates (Fig. S10a). These results do not support the earlier hypothesis that omnivory acts as a macroevolutionary sink across higher vertebrate taxa (Burin et al. 2016), instead suggesting that omnivory is a source of diversity. This demonstrates that a lack of standardization in classification can lead to vastly different conclusions regarding the macroevolutionary importance of a trait.

The species assigned to herbivore, carnivore, and omnivore differ between the Replicated and Price categorization methods (Table S3). Specifically, there are fewer herbivores and more omnivores when using EltonTraits data compared to when using the Price categorization. Different methodologies for measuring diet, such as analyses of stomach contents or direct behavioral observation, use different proxies as evidence of food item consumption and may ignore other evidence, leading to varying estimations of diet (e.g., McInnis et al. 1983; Shrestha and Wegge 2006; Balestrieri et al. 2011). Moreover, different approaches record diet across varying temporal scales, leading to incongruences in diet reconstruction for different species if a dataset uses multiple proxies to obtain its dietary data (Davis and Pineda-Munoz 2016). Use of different and varied sources of diet data by Wilman et al. (2014) and Price et al. (2012) likely compounded these incongruences, leading to disagreement regarding each species’ diet and subsequently causing discrepancies between the results of our diversification analyses when using the different datasets.

These results highlight the limitation of comparing studies that use different trait categorization methods and are applicable to other macroevolutionary studies where biologically continuous traits are discretized for analyses. Multiple complex, biologically continuous traits have been treated as qualitative variables in analyses of trait-dependent diversification, including morphology (e.g., Fernández-Mazuecos et al. 2013; Conway and Olsen 2019), locomotory mode (e.g., Nations et al. 2020), and biome or habitat usage (e.g., Schwery et al. 2015; Heinicke et al. 2017; Menéndez et al. 2020). However, methods exist to capture the continuous nature of these complex traits, so their macroevolutionary contributions can be assessed without the introduction of biases associated with variable discretization. The feasibility of such analyses has already been demonstrated for studies of morphology (Onstein et al. 2016) and habitat usage (Burbrink et al. 2012; Velasco et al. 2016). Locomotory mode can be predicted by morphology (Petrović et al. 2017; Verde Arregoitia et al. 2017), which is itself a continuous trait, meaning that locomotory mode may also be treated as biologically continuous in analyses of trait-dependent diversification. Given our findings, we recommend that future studies of trait-dependent speciation and diversification consider developing methods to capture the continuous nature of their trait of interest.

## Supporting Information

The doi for our data is 10.5061/dryad.bzkh1899n

