## Supporting Information for "Evolution of degrees of carnivory and dietary specialization across Mammalia and their effects on speciation"

### Supplementary Tables

**Table S1. The number of regime shifts in degree of carnivory and dietary specialization across Mammalia**

| Clade | Degree of carnivory | Degree of dietary specialization |
|---|---|---|
| | Number of shifts (Degree of Carnivory) | Number of shifts (Degree of Dietary Specialization) |
| Mammalia | 134 | 50 |
| Artiodactyla + Perissodactyla | 2 | 2 |
| Atlantogenata | 5 | 1 |
| Carnivora + Pholidota | 17 | 12 |
| Chiroptera | 13 | 2 |
| Euarchonta | 14 | 7 |
| Eulipotyphla | 2 | 3 |
| Lagomorpha | 0 | 0 |
| Marsupialia | 16 | 5 |
| Rodentia | 61 | 16 |

**Table S2. Selection strength (α) and phylogenetic half-life ($t_{1/2}$) for multi optimum Ornstein-Uhlenbeck (OU) models of trait evolution for carnivory and dietary specialization in Mammalia.** α, selection strength; $Rt_{1/2}$, relative phylogenetic half-life (represented as percentage of tree height); $At_{1/2}$, absolute phylogenetic half-life in Myr

| Carnivory | | | Dietary Specialization | | |
|---|---|---|---|---|---|
| α | $Rt_{1/2}$ (%) | $At_{1/2}$ (Myr) | α | $Rt_{1/2}$ (%) | $At_{1/2}$ (Myr) |
| 84.28 | 0.82 | 2.03 | 23.04 | 3.01 | 7.44 |

**Table S3. Contingency table of categorical diet classifications from Price et al. (2012) and this study.** The diet groupings of Price et al. (2012) were reused and pruned to species included in this study (Price categorization). The classification method of Price et al. (2012) was replicated using EltonTraits data, where 0% animal matter = herbivore, 1-99% = omnivore, and 100% = carnivore (Strict categorization).

| | Price | | |
|---|---|---|---|
| Strict | Herbivore | Omnivore | Carnivore |
| Herbivore | 441 | 63 | 1 |
| Omnivore | 134 | 250 | 35 |
| Carnivore | 1 | 34 | 364 |

#### Supplementary Figures

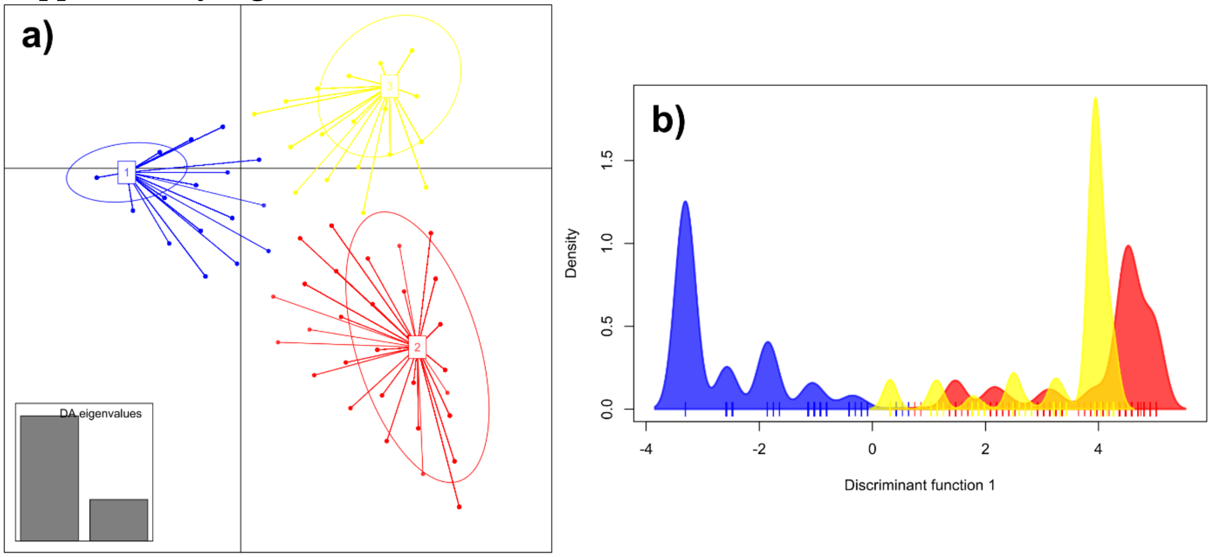

**Figure S1.** DAPC plot for construction of the degree of carnivory measure. Blue cluster = herbivore, red = vertivore, yellow = invertivore. a) Distribution of clusters on first and second discriminant functions. b) Distribution of herbivores, vertivores, and invertivores along the first discriminant function, which was later rescaled to produce the degree of carnivory variable.

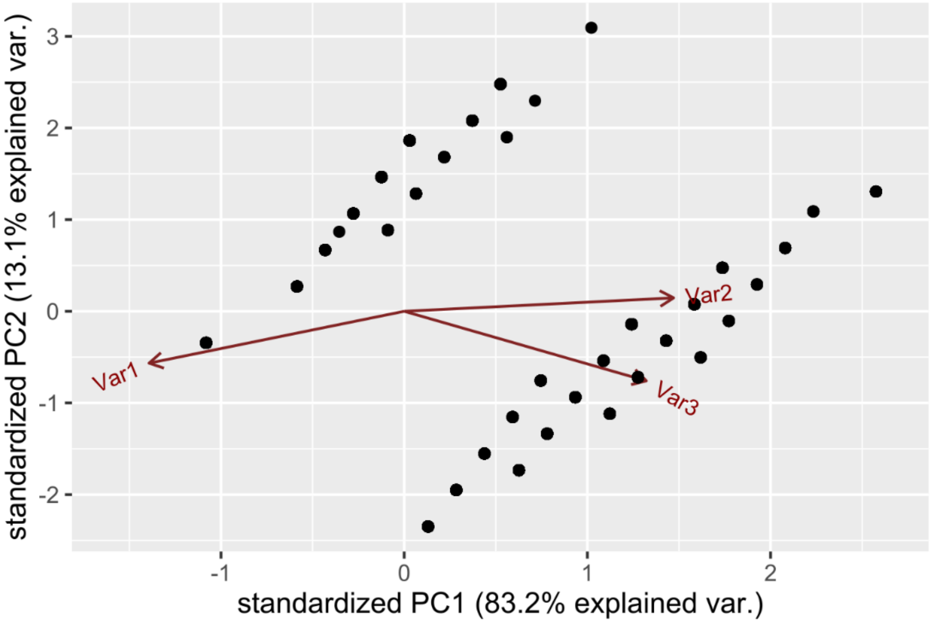

**Figure S2.** PCA of variables used to construct degree of dietary specialization measure. Scores on PC1 were later rescaled to produce the final dietary specialization variable. Var1 = percentage (from 20 – 100%) of a species' most common food type. Var2 = number of food types (from 1 – 6) included in each species' diet. Var3 = trophic breadth: species that consumed only plants or animals had a score of 1, those that consumed both food types had a score of 2.

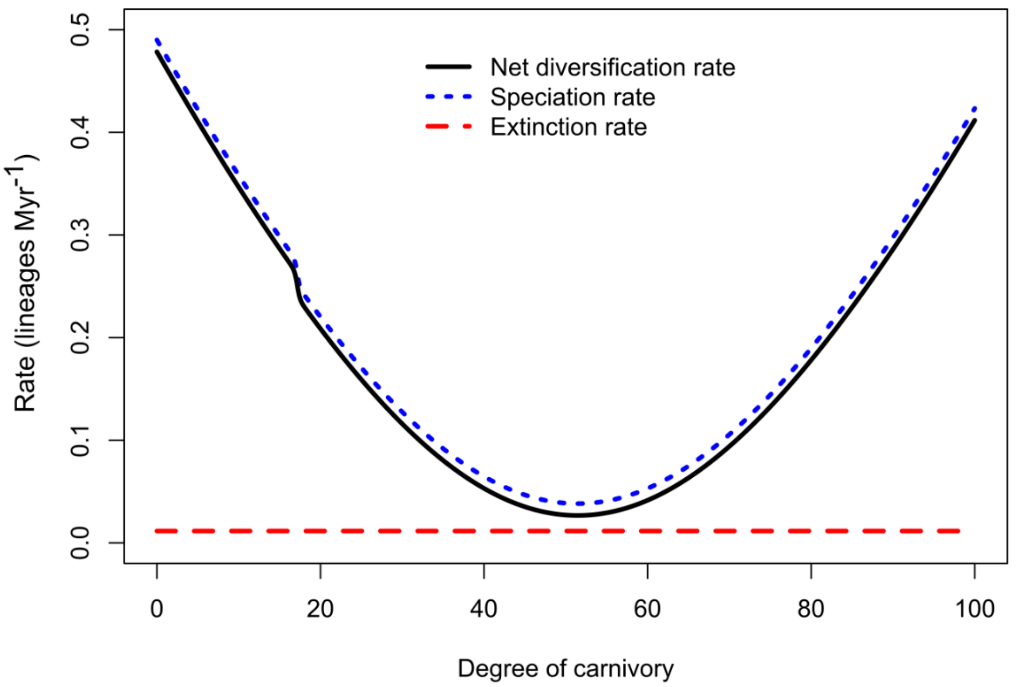

**Figure S3.** The relationship between degree of carnivory and speciation, extinction, and net diversification rates in Mammalia. The speciation, extinction, and net diversification rate lines represent the relationship produced by averaging over the significant QuaSSE models associated with the eleven phylogenies included in this study.

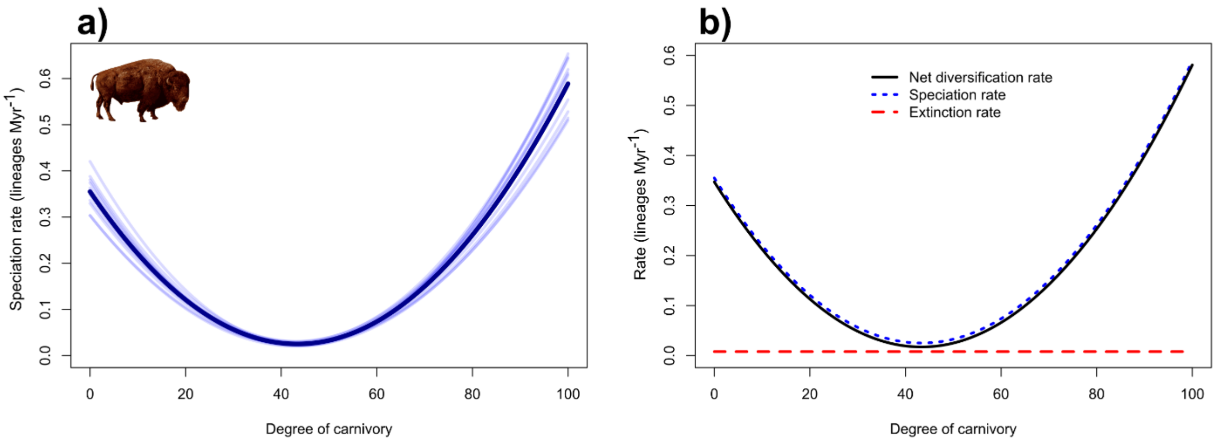

**Figure S4.** The relationship between degree of carnivory and speciation, extinction, and net diversification rate in Artiodactyla + Perissodactyla. Across the eleven trees used in this study, eleven were associated with a statistically significant best-fitting QuaSSE model in Artiodactyla + Perissodactyla. We consider this dynamic very robust to phylogenetic uncertainty. a) The transparent blue lines show the relationships between degree of carnivory and speciation rate predicted by these models. The dark blue line represents the relationship produced by averaging over each significant model. b) The speciation, extinction, and net diversification rate lines represent the relationship produced by averaging over the significant QuaSSE models. Image source: Wikimedia Commons.

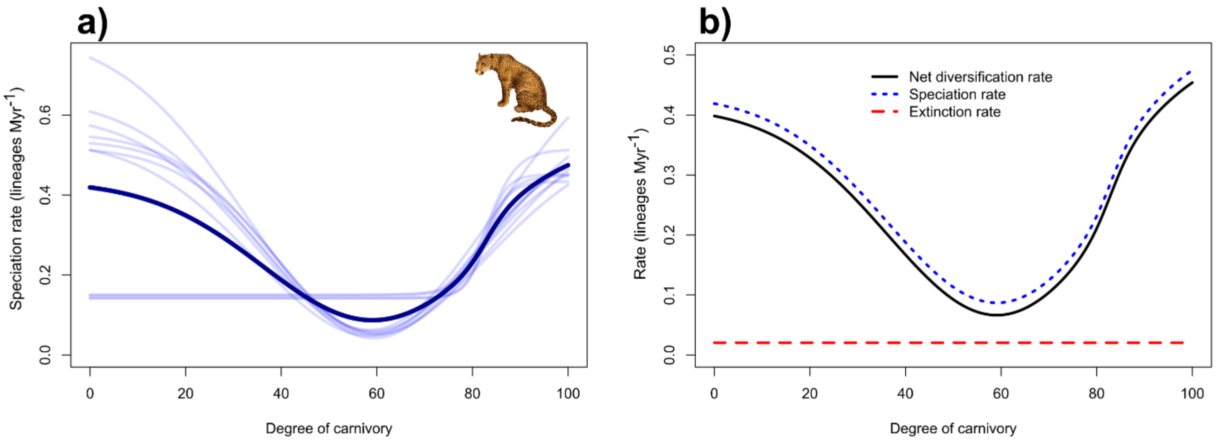

**Figure S5.** The relationship between degree of carnivory and speciation, extinction, and net diversification rate in Carnivora + Pholidota. Across the eleven trees used in this study, eleven were associated with a statistically significant best-fitting QuaSSE model in Carnivora + Pholidota. We consider this dynamic very robust to phylogenetic uncertainty. a) The transparent blue lines show the relationships between degree of carnivory and speciation rate predicted by these models. The dark blue line represents the relationship produced by averaging over each significant model. b) The speciation, extinction, and net diversification rate lines represent the relationship produced by averaging over the significant QuaSSE models. Image source: Wikimedia Commons.

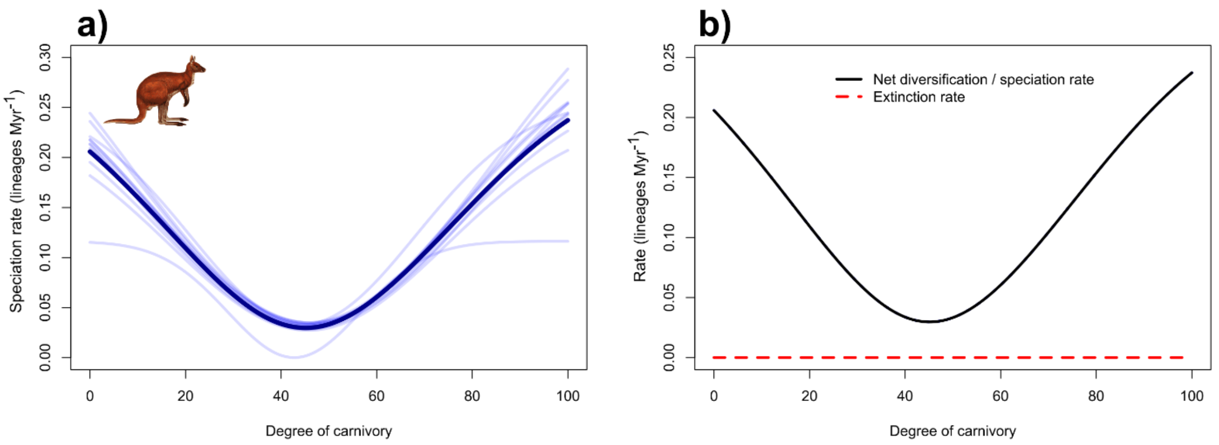

**Figure S6.** The relationship between degree of carnivory and speciation, extinction, and net diversification rate in Marsupialia. Across the eleven trees used in this study, eleven were associated with a statistically significant best-fitting QuaSSE model in Marsupialia. We consider this dynamic very robust to phylogenetic uncertainty. a) The transparent blue lines show the relationships between degree of carnivory and speciation rate predicted by these models. The dark blue line represents the relationship produced by averaging over each significant model. b) The extinction and net diversification rate lines represent the relationship produced by averaging over the significant QuaSSE models. Extinction rate was at a constant low level; thus, the speciation and net diversification rates were indistinguishable. Image source: Wikimedia Commons.

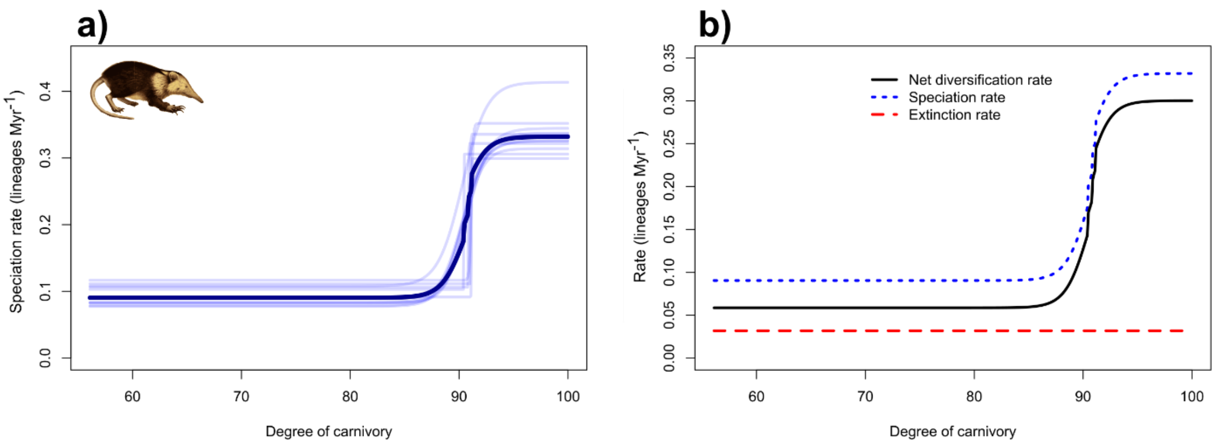

**Figure S7.** The relationship between degree of carnivory and speciation, extinction, and net diversification rate in Eulipotyphla. Across the eleven trees used in this study, nine were associated with a statistically significant best-fitting QuaSSE model in Eulipotyphla. We consider this dynamic only moderately robust to phylogenetic uncertainty. a) The transparent blue lines show the relationships between degree of carnivory and speciation rate predicted by these models. The dark blue line represents the relationship produced by averaging over each significant model. b) The speciation, extinction, and net diversification rate lines represent the relationship produced by averaging over the significant QuaSSE models. Image source: Wikimedia Commons.

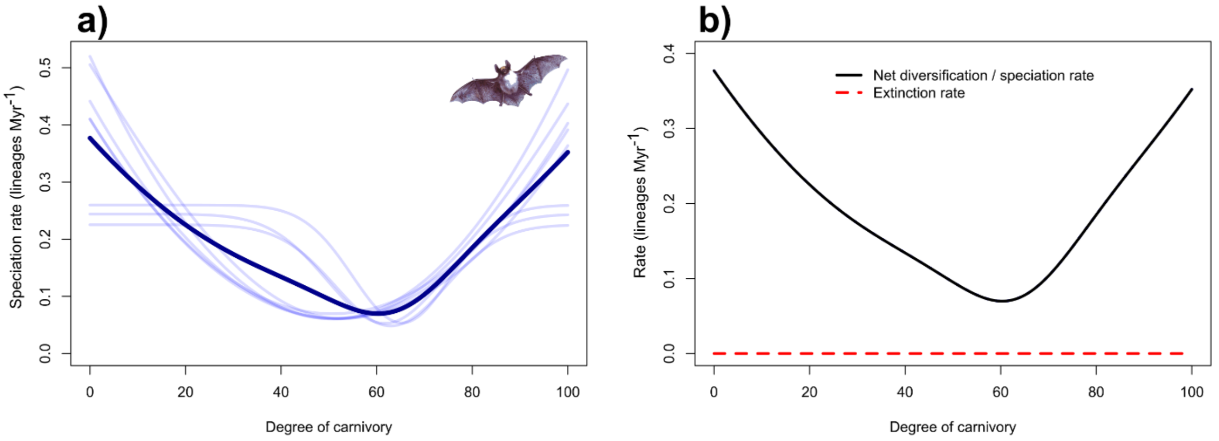

**Figure S8.** The relationship between degree of carnivory and speciation, extinction, and net diversification rate in Chiroptera. Across the eleven trees used in this study, eight were associated with a statistically significant best-fitting QuaSSE model in Chiroptera. We consider this dynamic only moderately robust to phylogenetic uncertainty. a) The transparent blue lines show the relationships between degree of carnivory and speciation rate predicted by these models. The dark blue line represents the relationship produced by averaging over each significant model. b) The extinction and net diversification rate lines represent the relationship produced by averaging over the significant QuaSSE models. Extinction rate was at a constant low level; thus, the speciation and net diversification rates were indistinguishable. Image source: Wikimedia Commons.

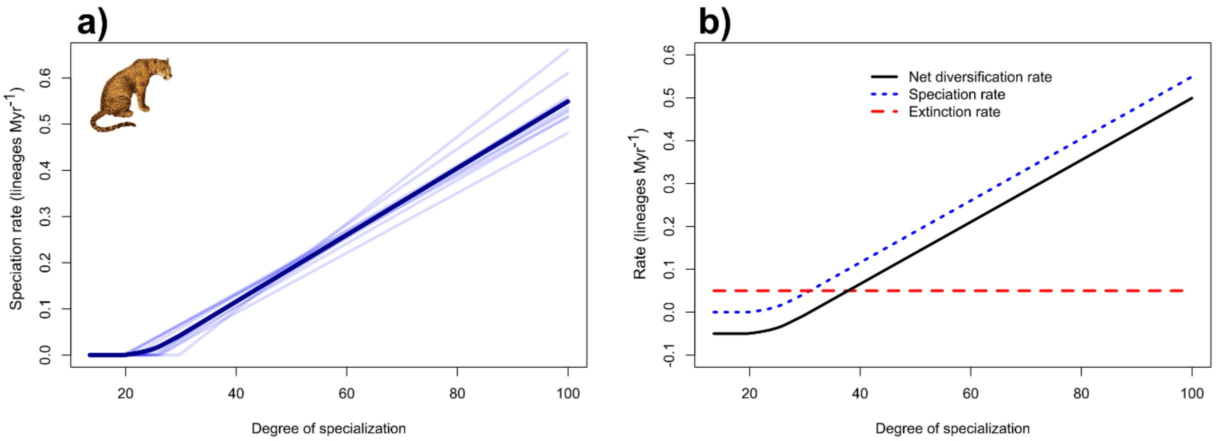

**Figure S9.** The relationship between degree of dietary specialization and speciation, extinction, and net diversification rate in Carnivora. Across the eleven trees used in this study, eleven were associated with a statistically significant best-fitting QuaSSE model in Carnivora. We consider this dynamic very robust to phylogenetic uncertainty. a) The transparent blue lines show the relationships between degree of dietary specialization and speciation rate predicted by these models. The dark blue line represents the relationship produced by averaging over each significant model. b) The speciation, extinction, and net diversification rate lines represent the relationship produced by averaging over the significant QuaSSE models. Image source: Wikimedia Commons.

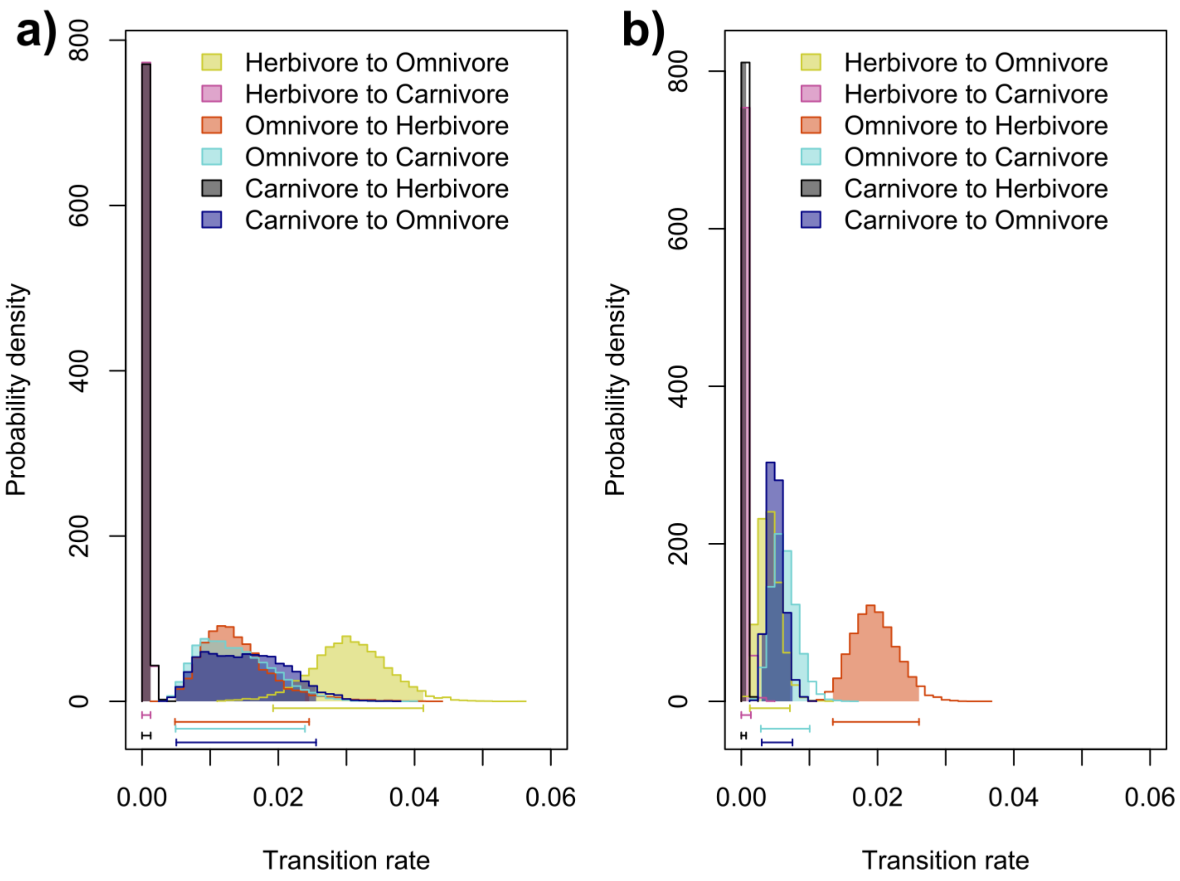

**Figure S10.** Posterior probability densities for transition rates between herbivores, omnivores, and carnivores, using three diet classification schemes. a) The diet groupings of Price et al. (2012) were reused with the most recent mammalian phylogeny (Upham et al. 2019). b) The classification method of Price et al. (2012) was replicated using data from EltonTraits.

**Dietary regimes in Carnivora + Pholidota**

At its origin, members of Carnivora still occupied the original carnivory regime present at the root of Mammalia. This original regime was one of moderately high carnivory, with an optimum score indicating a diet comprising 80-90% animal matter (Fig. S11). Pholidota also occupies this regime. Contemporary carnivoran families with members that still occupy this original regime include Mephitidae, Canidae, Viverridae, Herpestidae, Eupleridae, and Hyaenidae. A shift towards a similar carnivory regime occurred at the origin of Mustelidae and is still occupied by members of Helictidinae, Guloninae, Mellivorinae, and Taxidiinae. Four independent shifts towards hypercarnivory occurred in Carnivora, in: Felidae, Pinnipedia, the branch leading to Mustelinae, Ictonychinae, and Lutrinae, and the branch leading to *Ursus maritimus*. Shifts towards varying degrees of herbivory occurred throughout Carnivora. A shift to mesoherbivory occurred in Musteloidea and is still occupied by *Ailurus fulgens* and members of Procyonidae. Similar shifts occurred in Nandiniidae and in Genettinae at the branch leading to *Poiana richardsonii*. There was a shift to hyperherbivory at the origin of Ursidae, followed by a shift to omnivory at the origin of Ursinae, then a subsequent shift to mesoherbivory at the branch leading to *Ursus*. Other shifts to omnivory occurred in Melinae, *Procyon*, Paradoxurinae, *Urocyon*, and the branch leading to *Vulpes cana*.

For dietary specialization, a regime shift towards a more generalized optimum diet occurred at the origin of Carnivora (Fig. S11). This optimum specialization score is similar to diets comprising two food types, one animal-based and one plant-based, with a bias towards one of the two food types. Most members of Carnivora still occupy this original specialization regime. Pholidota occupies the older, more specialized regime. Shifts towards greater dietary specialization occurred in Genettinae, Hemigalinae, the branch leading to *Ursus maritimus*, and the branch leading to Mustelinae, Ictonychinae, and Lutrinae. Multiple shifts to higher specialization occurred throughout Canidae. A shift to a decreased optimum degree of dietary specialization occurred at the branch leading to *Mustela itatsi*.

*Discussion*

Mesocarnivory has previously been reported as the basal condition in Carnivora (Van Valkenburgh 2007; Slater and Friscia 2019). However, our study reported a high optimal degree of carnivory at the origin of Carnivora, suggesting disagreement with previous findings. Earlier studies only considered the proportion of vertebrates in a given species' diet when measuring carnivory (Van Valkenburgh 1988, 2007), whereas we also incorporated invertivory into our carnivory variable. Thus, invertivorous species are considered to have higher degrees of carnivory in this study than they were previously. Extant species with similar carnivory scores to the original optimum value found in this study are predominantly invertivorous and would be considered mesocarnivorous by earlier metrics. Therefore, the optimum degree of carnivory found at the origin of Carnivora supports the findings of earlier studies.

The placement of ecomorphological shifts within Carnivora by Slater and Friscia (2019) is in loose agreement with the results of our study. Whereas the ancestral carnivory regime in our analysis extends to include Mephitidae, the earlier study instead reports a shift to omnivory near the base of Caniformia (excluding Canidae). Also, fewer shifts within Feliformia occur in our results. Additional shifts at lower taxonomic levels occur in our study – for example within Ursidae and Canidae – which do not occur in Slater and Friscia (2019).

Carnivora + Pholidota was the only subclade in our analysis to return a significant trait-dependent speciation dynamic for both degree of carnivory and degree of dietary specialization. Carnivoran speciation favored increased specialization for either end of the carnivory spectrum. An evolutionary trend of increasing body size over time, known as Cope's Rule, is documented in Carnivora, as well as in other mammalian lineages (Alroy 1998). Within Carnivora, there is an association between increasing body size and specialization for hypercarnivory (Van Valkenburgh et al. 2004). This, in combination with the high energy content and easy digestibility of a carnivorous diet, has been posited as the reason for a bias towards evolution of hypercarnivorous feeding strategies in the carnivoran fossil record (Van Valkenburgh 2007). Increased vulnerability to extinction and reduced species duration of hypercarnivorous lineages has resulted in the repeated appearance of convergent carnivorous forms in distantly related species throughout Carnivora (Van Valkenburgh 1991, 2007; Balisi et al. 2018). This evolutionary trend favoring specialization for hypercarnivory explains the positive relationship between increasing degrees of carnivory and speciation rate observed for Carnivora in our analyses.

A positive relationship between hyperherbivory and speciation rate is surprising and is not explained using the same trend. This relationship may instead be driven by the dietary specialization-dependent speciation dynamic observed in Carnivora. Constraints may be placed on the evolution of herbivorous carnivorans due necessarily to their descent from carnivorous ancestors (Figueirido et al. 2010), which can explain why shifts to herbivory are infrequent in the order despite the potential for herbivores to speciate quickly.

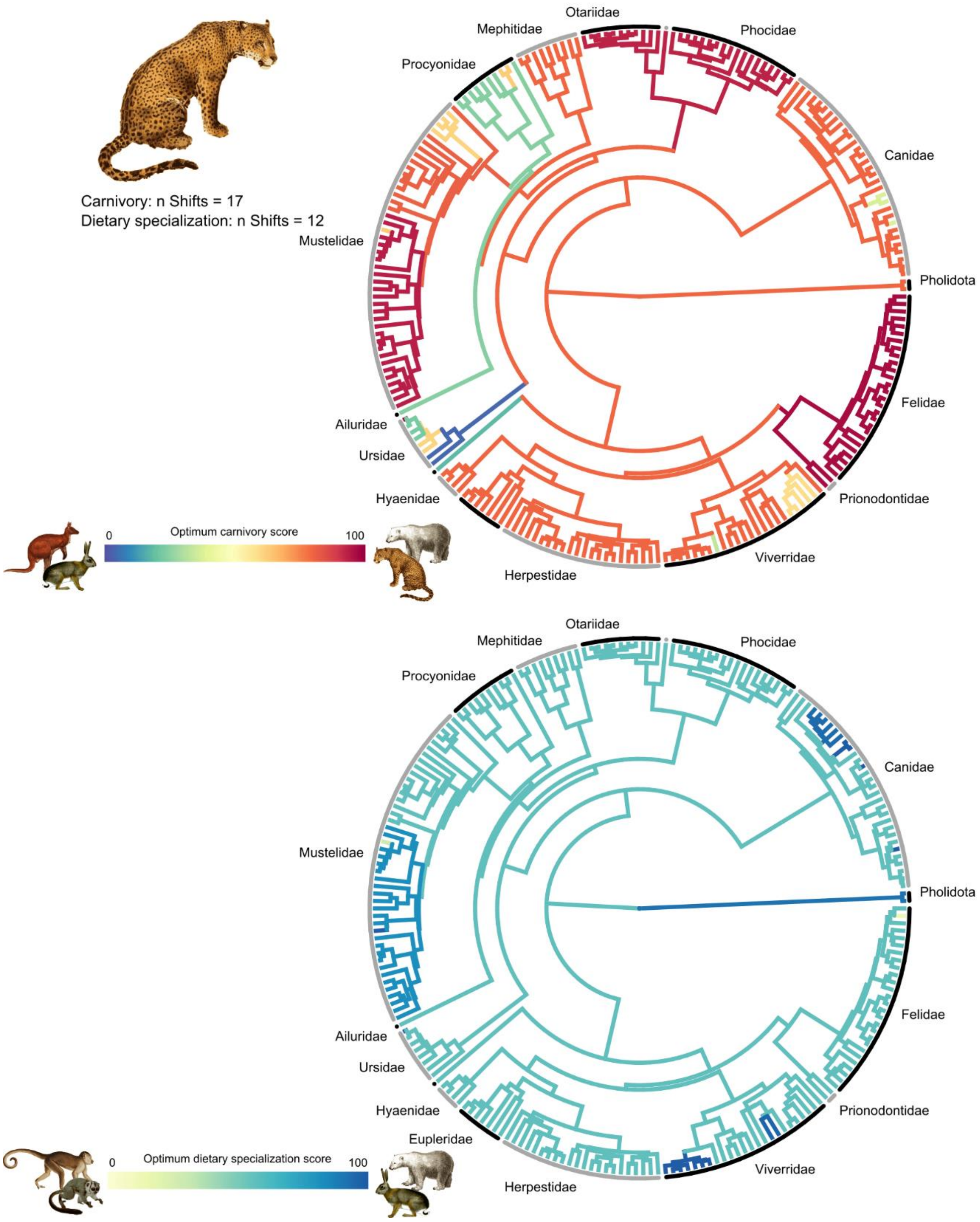

**Figure S11.** Regime shifts in optimum carnivory score (top) and optimum dietary specialization score (bottom) in Carnivora + Pholidota. Branch color indicates the value of the optimum diet score. Position of color changes represents the location of regime shifts**.** Images source: Wikimedia Commons.

**Dietary regimes in Artiodactyla + Perissodactyla**

The optimum carnivory score of the regime occupied at the origin of Artiodactyla + Perissodactyla was zero, representing a completely herbivorous diet (Fig. S12). This hyperherbivorous regime is still occupied by all perissodactyls and most artiodactyls. A shift towards hypercarnivory occurred in the branch leading to Cetacea. A shift to mesoherbivory occurred at the branch leading to *Sus* and *Potamochoerus* within Suidae.

The optimum degree of dietary specialization at the origin of Artiodactyla + Perissodactyla was high, representing specialization on a single food type (Fig. S12). Shifts towards dietary generalism occurred at the origins of Cephalophinae and Suina.

*Discussion*

Very few shifts in degree of carnivory or degree of dietary specialization occurred in Artiodactyla. Most artiodactylans are hyperherbivorous, with cetaceans representing a single clade of hypercarnivores. Although there is little evidence for a rapid adaptive radiation within Cetacea (Steeman et al. 2009; Slater et al. 2010), extant whales represent a relatively speciose clade compared to some other ungulate groups. This is especially true in comparison to clades with omnivorous members, which are restricted to the smaller Suina suborder. This could explain the significant relationship between degree of carnivory and speciation rate observed for Artiodactyla + Perissodactyla, where herbivores and carnivores have higher speciation rates than more omnivorous species.

There was no significant relationship between degree of dietary specialization and speciation rate in Artiodactyla + Perissodactyla. Generalist, mixed-feeders were previously shown to have higher diversification rates than browse-specialist ruminants (Cantalapiedra et al. 2014). However, specialist grazers also had higher diversification rates than browse-specialists, indicating that there is not a strictly dietary specialization-dependent diversification dynamic occurring within Artiodactyla. Additionally, browsers, grazers, and mixed-feeders are all scored as folivore specialists in our study, meaning that this diversification dynamic cannot be captured using our variable. Studies assessing diet at different levels of coarseness detect different relationships between diet and diversification. These studies are complementary to one another and it is important to continue assessing at different scales to elucidate the full picture of diet-dependent diversification across different taxa.

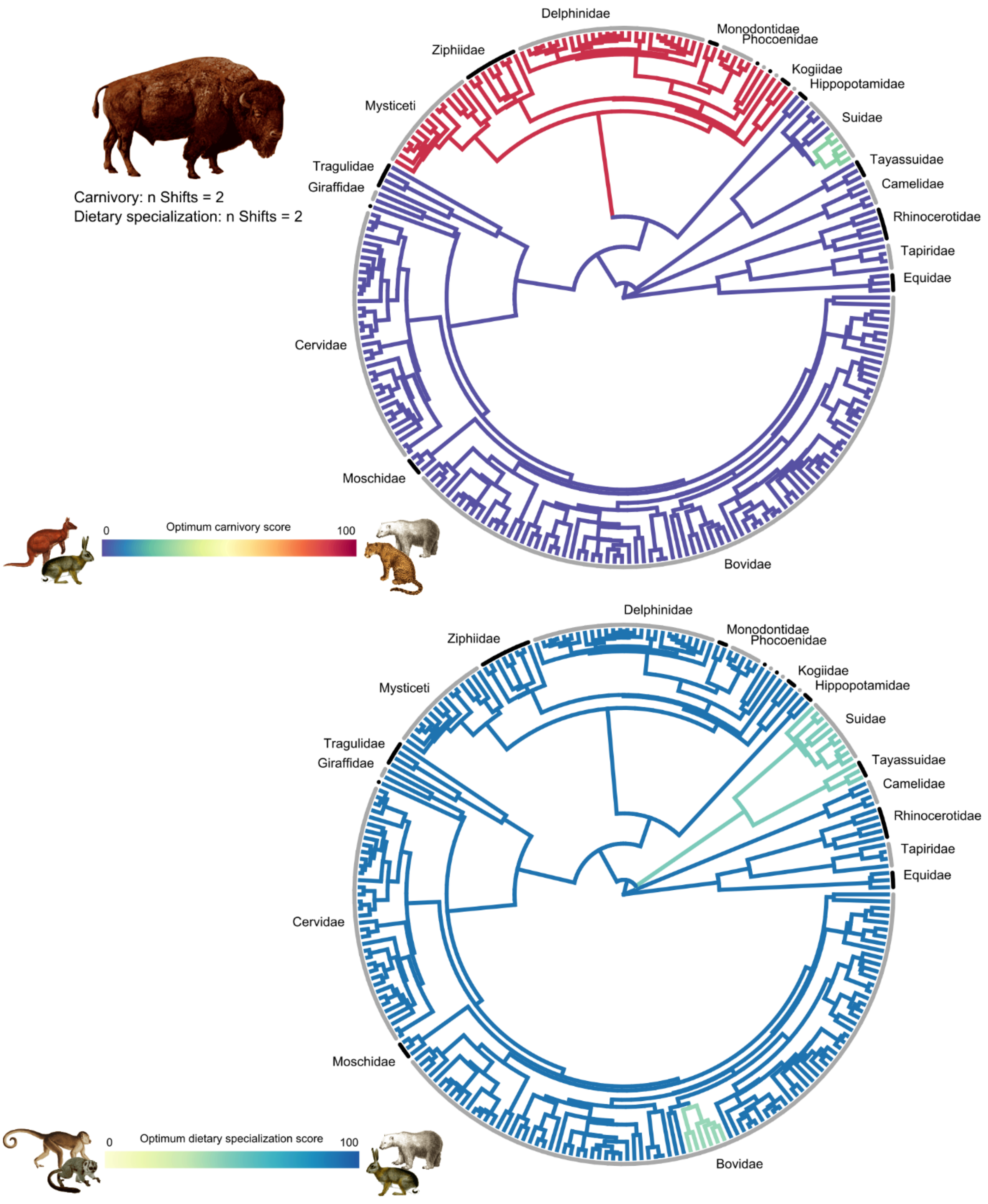

**Figure S12.** Regime shifts in optimum carnivory score (top) and optimum dietary specialization score (bottom) in Artiodactyla + Perissodactyla. Branch color indicates the value of the optimum diet score. Position of color changes represents the location of regime shifts. Images source: Wikimedia Commons.

**Dietary regimes in Chiroptera**

At the origin of Chiroptera, a carnivory regime arose with an optimum carnivory score similar to extant species whose diets are comprised exclusively of invertebrates (Fig. S13). This optimum score is slightly higher than that of the original regime occupied at the root of Mammalia, which Chiroptera shifted from. All chiropteran families occupy this original regime, excluding Pteropodidae and Phyllostomidae. Most regime shifts in degree of carnivory and all shifts in dietary specialization occured in Phyllostomidae; shifts were infrequent throughout the rest of Chiroptera. At the origin of Phyllostomidae, there was a shift to a carnivory regime with an optimal omnivorous diet comprising equal parts invertebrates and plant matter.

Regime shifts to an optimum carnivory value representing a hyperherbivorous diet occurred at the origin of Pteropodidae and throughout Phyllostomidae, specifically at the origin of Stenodermatinae and the branches leading to the genera *Carollia*, *Phylloderma*, and *Platalina*. Mesoherbivorous regimes arose at the origin of Glossophaginae, the origin of *Ariteus*, and in the branch leading to *Phyllostomus discolor*. Shifts to mesocarnivorous regimes occurred at the origin of Phyllostominae and in the branch leading to *Mimon bennettii* and Glyphonycterinae. One shift to a regime consistent with a vertivorous optimum diet occurred at the origin of Desmodontinae.

Most of Chiroptera occupies the original dietary specialization regime occupied by the root of Mammalia and shared by most contemporary mammalian clades (Fig. S13). This optimum diet is highly specialized. The only shifts in dietary specialization occurred in Phyllostomidae. At the origin of the family, there was a shift to an optimum degree of dietary specialization similar to extant species that do not eat both plant and animal matter, but do consume multiple different foodstuffs. At the origin of Glossophaginae, there was further shift to a more generalized optimum, similar to extant diets comprising both animal and plant matter, but with a bias towards one over the other.

*Discussion*

Invertivory was the basal condition in Chiroptera, reinforcing the consensus that the common ancestor was insectivorous (Gunnell and Simmons 2005). Our results show that the optimal diet at the origin of Phyllostomidae – the clade with most dietary diversity (Freeman 2000) – was omnivorous. Previous findings reported that the ancestral diet of Phyllostomidae was insectivorous (Datzmann et al. 2010). However, our study indicated that regime shifts in degrees of carnivory and dietary specialization occurred at the origin of the family. These shifts were associated with an optimal diet that showed a decrease in dietary specialization in favor of increasing herbivory. Our findings align with recent studies that identified molecular adaptations for processing diverse macronutrients (Potter et al. 2021) and development of an omnivorous sensory morphotype (Hall et al. 2021) on the branch leading to Phyllostomidae. These results suggest that the ancestral phyllostomid diet was relatively generalized and that the foundation for phyllostomid dietary diversity was laid at its origin.

We observed a moderately robust relationship between speciation rate and degree of carnivory in Chiroptera. Although the majority of extant bats are reported to be united by a single diversification process (Shi and Rabosky 2015), shifts in diversification rate have been inferred within chiropteran clades (Jones et al. 2005) and some of these have been attributed to changes in feeding strategy (Dumont et al. 2011; Monteiro and Nogueira 2011). We found that speciation rates are highest at either extreme end of the carnivory score spectrum, with omnivorous and mesocarnivorous species having the lowest rates (Fig. S6). Carnivorous diets predominate in Chiroptera and hyperherbivory is prevalent in two diverse chiropteran clades – Phyllostomidae and Pteropodidae. Conversely, omnivorous and mesocarnivorous chiropterans are comparatively rare, existing in small numbers primarily within Phyllostomidae. Thus, the distribution of carnivory scores across Chiroptera is consistent with the relationship between degree of carnivory and speciation rate that we detected.

We did not observe a significant relationship between degree of dietary specialization and speciation rate in Chiroptera. This result is perhaps unsurprising, given that earlier studies of dietary specialization's impact on chiropteran diversification have reached contradicting conclusions. Specialization for frugivory has been implicated as a cause for increased diversification in Neotropical noctilionoid bats (Rojas et al. 2012). However, a more recent study of noctilionoid bats reported that generalist herbivores speciate faster than specialized herbivores (Rojas et al. 2018). Our study examined the taxonomic breadth of Chiroptera and assessed the impact of dietary specialization independently of degree of carnivory. So, dietary specialization-dependent speciation dynamics may exist within smaller subsets of Chiroptera, but these relationships are not representative of all bats or diet types.

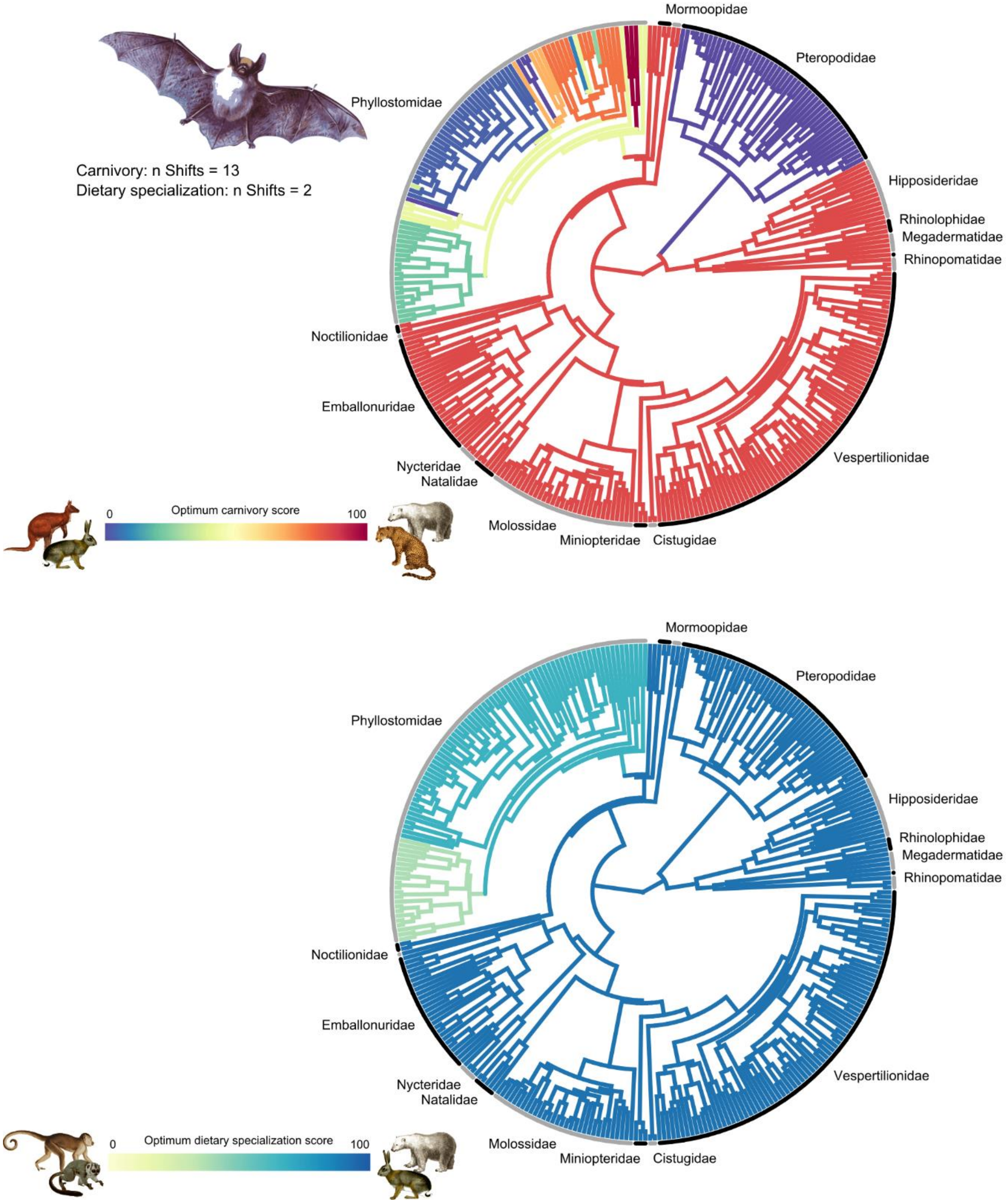

**Figure S13.** Regime shifts in optimum carnivory score (top) and optimum dietary specialization score (bottom) in Chiroptera. Branch color indicates the value of the optimum diet score. Position of color changes represents the location of regime shifts. Images source: Wikimedia Commons.

**Dietary regimes in Euarchonta**

At the origin of Euarchonta, the optimum carnivory score was one of mesoherbivory (Fig. S14). Only Primates remained in this regime, as the origins of Dermoptera and Scandentia coincide with shifts to new regimes with complete herbivory and high carnivory as optimum diets, respectively. Contemporary members of the original euarchontan carnivory regime still exist within Strepsirrhini, specifically within Daubentoniidae, Galagidae, and Lorisidae. There was a shift to complete herbivory on the branch leading to Cheirogalidae, Indriidae, Lemuridae, and Lepilemuridae. Shifts to mesocarnivorous optimum diets occurred at the origin of Lorisinae, *Paragalago,* and *Galagoides*. Outside of Strepsirrhini, there was a shift to high herbivory at the origin of Simiiformes, followed by a subsequent shift to complete herbivory at the origin of Colobinae. A shift to an optimum diet of mesoherbivory occurred in the branch leading to Cebidae, Callitrichidae, and Aotidae. There were four shifts to omnivorous optimum diets in Primates: in the branch leading to *Macaca fascicularis*, the branch leading to *Papio anubis*, the branch leading to *Allenopithecus* and *Miopithecus*, and the branch leading to *Mirza* and *Microcebus*. There was one shift to an optimal diet of complete carnivory at the origin of Tarsiidae.

Euarchonta occupied the same high-scoring dietary specialization regime as occupied at the root of Mammalia (Fig. S14). However, only contemporary members of Dermoptera and Scandentia still occupy this regime within Euarchonta. At the origin of Primates, there was a shift to a lower optimum dietary specialization score consistent with consuming three different food types, but with a bias towards one foodstuff. This regime is still occupied by extant members of Strepsirrhini and Tarsiidae. At the origin of Simiiformes, there was a subsequent shift with a decrease in optimum dietary specialization score, this time to an optimum diet comprising both plant and animal matter, but still with a bias to one over the other. A similar regime arose on the branch leading to *Ptilocercus* in Scandentia. At the origin of Colobinae, there was a shift back to a higher degree of dietary specialization, similar to the regime at the origin of Primates. There was a single shift to a highly generalized optimum diet consistent with consuming four to five different food types in the branch leading to *Mirza* and *Microcebus*.

*Discussion*

It has been suggested that the origin of Primates involved an adaptive shift towards a new feeding strategy, with competing hypotheses that this shift either represented a transition from ancestral insectivory to herbivory (Szalay 1968) or a transition to visual predation on insects in the lower canopy (Cartmill 1974). Our results instead indicate that the primate origin is associated with continued occupancy of the basal euarchontan carnivory regime, with an optimal diet of mesoherbivory. A previous finding that there was not an adaptive shift in chewing behavior at the origin of Primates was used to indicate that changes in food acquisition were decoupled from changes in jaw-muscle activity during primate evolution (Vinyard et al. 2007). However, those results could instead provide further evidence for the lack of a dietary shift at the origin of Primates.

The basal condition of Euarchonta is contended to be most similar to *Ptilocercus*, considered the most plesiomorphic extant tree shrew (Szalay and Drawhorn 1980). Thus, the ancestral euarchontan was suggested to be predominantly insectivorous like *Ptilocercus* (Bloch et al. 2007). This contradicts the results of our regime shift analyses, where a shift to a carnivorous optimum diet arose in Scandentia, from an ancestrally mesoherbivorous regime. Our finding of an herbivorous optimal diet at the origin of Euarchonta is supported by reconstruction of ancestral bitter taste receptor (TAS2R) gene numbers within Euarchontoglires (Hayakawa et al. 2014).

*Tas2r* gene number is positively correlated with the proportion of plant matter in a given species' diet (Li and Zhang 2013). At the origin of Euarchonta, *Tas2r* gene number was high, remained high throughout the evolution of Primates, but decreased with the divergence of the insectivorous Scandentia.

We found no significant relationship between speciation rate and degree of carnivory or degree of dietary specialization in Euarchonta. An earlier study found that primates possessing mutualistic dietary interactions with plants (i.e. frugivores) had higher diversification rates than those with antagonistic interactions (i.e. folivores and insectivores) (Gómez and Verdú 2012). Our measure of degree of carnivory does not differentiate between folivory and frugivory, so both forms of herbivory are attributed the same carnivory score. It would therefore not be possible to observe the same diet-dependent diversification dynamic using our study design. Thus, the results of the two studies do not compete against one another. However, a more recent study showed that insectivorous primate lineages have diversified more slowly than herbivorous lineages (Scott 2019). This diversification dynamic is driven by a low rate of diversification in nocturnal lineages relative to diurnal ones, as there is an association between insectivory and nocturnality in Primates. Reasons for our inability to return the same significant relationship between degree of carnivory and speciation rate may include our use of a continuous variable instead of categorical one, or by expansion of our trait-dependent speciation analyses to all euarchontans.

Lack of an observable relationship between dietary specialization and speciation rate in Euarchonta corroborates previous findings. Specialization for folivory was not related to changes in diversification rate in African colobine monkeys but was shown to depress diversification in Asian colobines (Tran 2014). Primate frugivory-specialist lineages did not show significantly different net diversification rates to non-frugivores (Scott 2018, but see Gómez and Verdú 2012). These results indicate that dietary specialization – measured at either a finer scale as in the previous studies or more broadly as in our study – does not have a significant impact on diversification rate in Euarchonta.

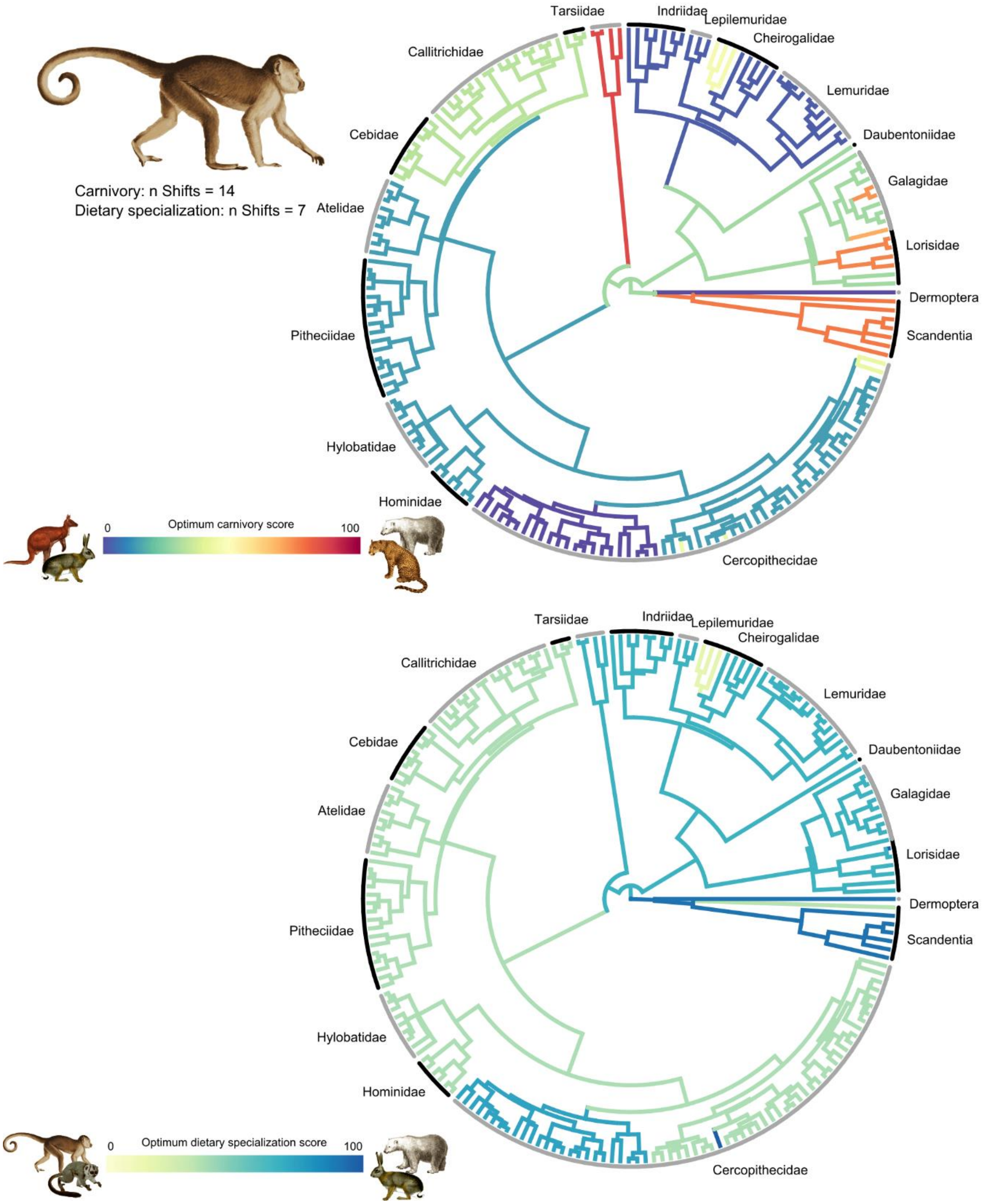

**Figure S14.** Regime shifts in optimum carnivory score (top) and optimum dietary specialization score (bottom) in Euarchonta. Branch color indicates the value of the optimum diet score. Position of color changes represents the location of regime shifts. Images source: Wikimedia Commons.

#### Dietary regimes in Eulipotyphla

Within Eulipotyphla, only Solenodontidae remains in the original carnivory regime occupied at the root of Mammalia (Fig. S15). In the branch leading to the rest of Eulipotyphla, there arose a shift towards an optimum carnivory score indicative of a completely carnivorous diet, with heavy bias towards invertivory over vertivory. This carnivory regime is occupied by almost all eulipotyphlans. A second shift in carnivory score occurred at the origin of *Blarina*, to a mesocarnivorous optimum diet.

Most of Eulipotyphla still occupies the dietary specialization regime found at the root of Mammalia (Fig. S15). Three shifts to reduced dietary specialization occurred in Eulipotyphla. At the origin of Solenodontidae, there was a shift to an optimum specialization score consistent with an omnivorous diet comprising four different food types. Within Erinaceinae (excluding *Erinaceus*), there was a shift to an omnivorous optimum diet comprising three different food types. In the branch leading to *Blarina* and *Sorex*, there was also a shift to an optimum diet comprising three food types.

*Discussion*

Eulipotyphla shows less dietary diversity than many other mammalian clades, as most members are predominantly invertivorous or vertivorous, with some rarer omnivorous members included. The evolution of eulipotyphlan diet is not well studied so there is little to compare our regime shift analyses to. Our results provide support for a previously suggested insectivorous origin for Eulipotyphla (Wu et al. 2017). A shift to increased diversification rate was previously reported in a branch leading to *Crocidura*, the most speciose genus within Eulipotyphla (Rossi et al. 2018). Members of *Crocidura* are coded as consuming equal proportions of invertebrates and vertebrates in the EltonTraits dataset (Wilman et al. 2014), which would increase their carnivory score relative to predominantly invertivorous eulipotyphlans. This could explain the association between extreme hypercarnivory and heightened speciation rate observed for Eulipotyphla in our study.

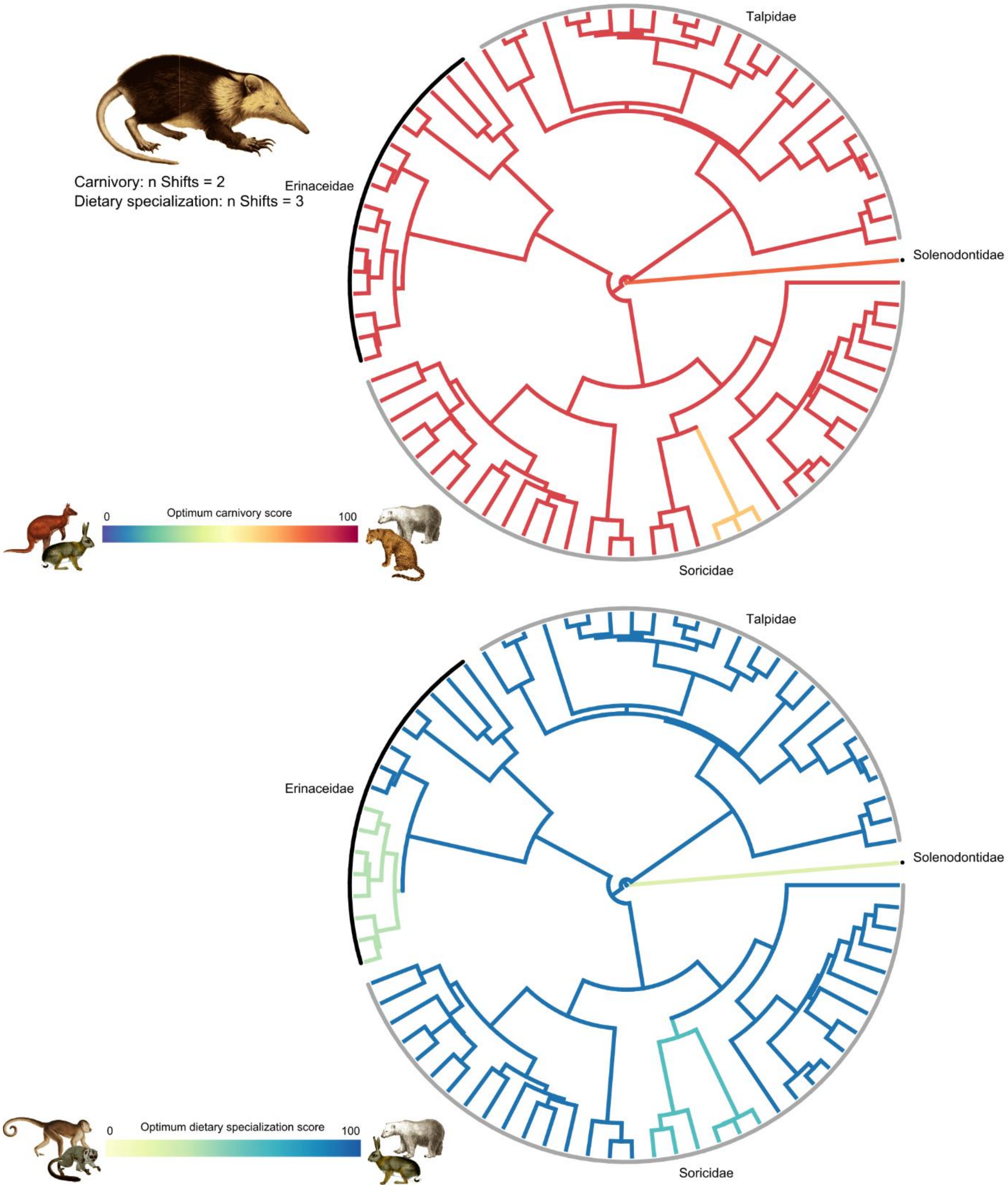

**Figure S15.** Regime shifts in optimum carnivory score (top) and optimum dietary specialization score (bottom) in Eulipotyphla. Branch color indicates the value of the optimum diet score. Position of color changes represents the location of regime shifts. Images source: Wikimedia Commons.

**Dietary regimes in Lagomorpha**

All members of Lagomorpha belong to a single dietary regime for both degree of carnivory and degree of dietary specialization (Fig. S16). The carnivory regime occupied by Lagomorpha arose at the origin of Glires and has an optimum carnivory score representing complete herbivory. The high-scoring dietary specialization regime occupied by Lagomorpha is the same as is occupied at the root of Mammalia.

**Dietary regimes in Atlantogenata**

The origin of Atlantogenata is associated with the same highly carnivorous carnivory regime occupied at the root of Mammalia (Fig. S17). This ancestral regime is still occupied by Orycteropodidae and members of Vermilingua and Cingulata in Xenarthra. In the branch leading to Afroinsectiphilia in Afrotheria, there was a regime shift to a carnivory score consistent with a completely invertivorous diet in extant species. Shifts to herbivorous optimum diets occurred at the origins of Folivora and Paenungulata. Shifts to omnivorous optimum diets occurred in the branch leading to *Echinops telfairi* and the branch leading to *Chaetophractus* and *Euphractus*.

Most of Atlantogenata occupies the same dietary specialization regime that arose at the root of Mammalia, and that is shared by most mammalian lineages (Fig. S17). One shift in dietary specialization occurred on the branch leading to *Chaetophractus* and *Euphractus*, leading to a moderately generalized optimum diet.

*Discussion*

Because the grouping of Afrotheria and Xenarthra into Atlantogenata has historically been contentious (Esselstyn et al. 2017; Upham et al. 2019), there is a dearth of literature on the evolution of diet within Atlantogenata. The ancestral condition of Xenarthra has been reconstructed as myrmecophagous due to the dramatically reduced dentition observed in xenarthrans, which is usually associated with invertivory (Gaudin and Croft 2015). This agrees with the results of our regime shift analyses. Our results also indicate the ancestral optimal condition of Afrotheria was one of high carnivory, with the common ancestor of Afrotheria sharing the same dietary regime as the common ancestor of Xenarthra.

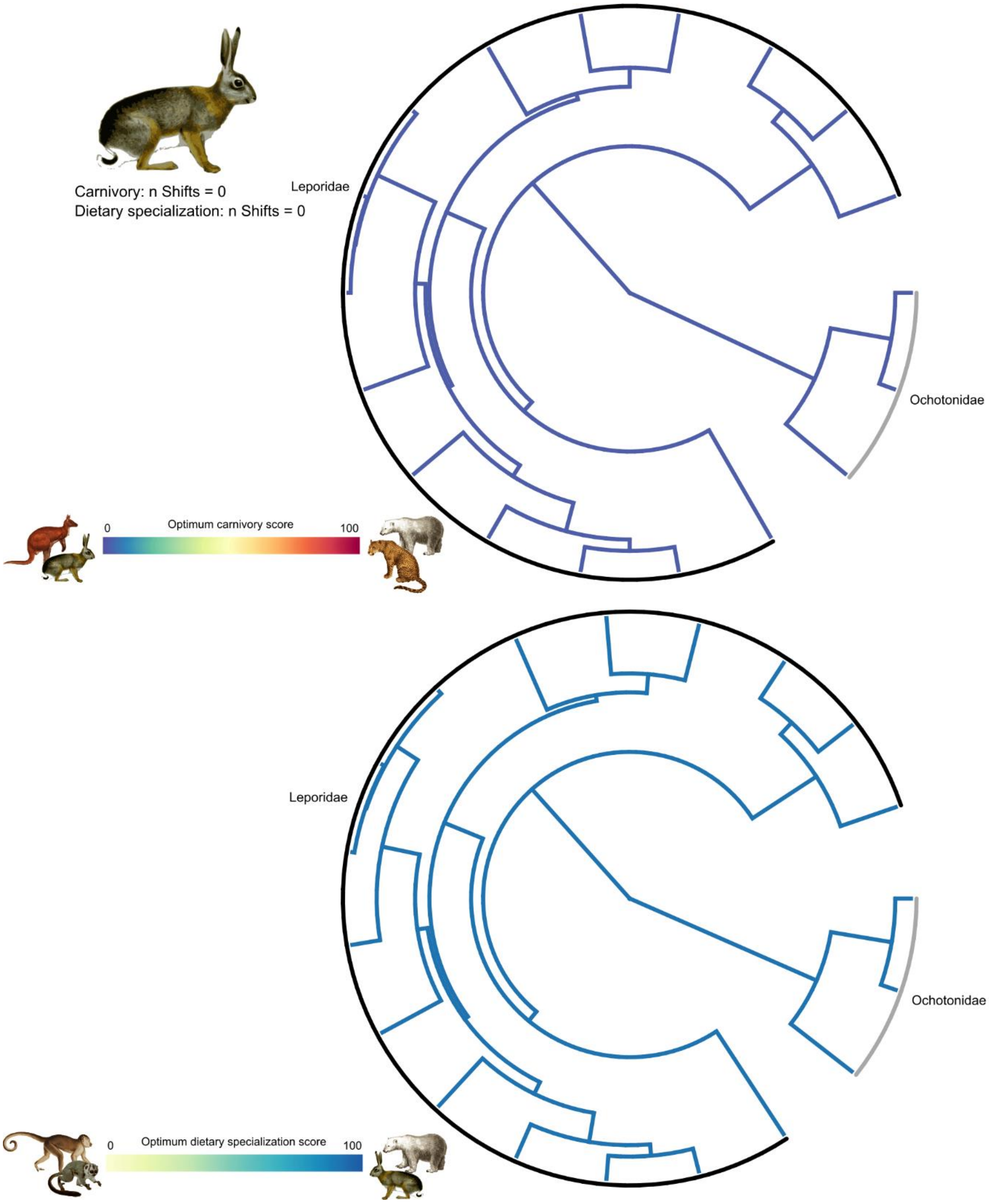

**Figure S16.** Regime shifts in optimum carnivory score (top) and optimum dietary specialization score (bottom) in Lagomorpha. Branch color indicates the value of the optimum diet score. Position of color changes represents the location of regime shifts. Images source: Wikimedia Commons.

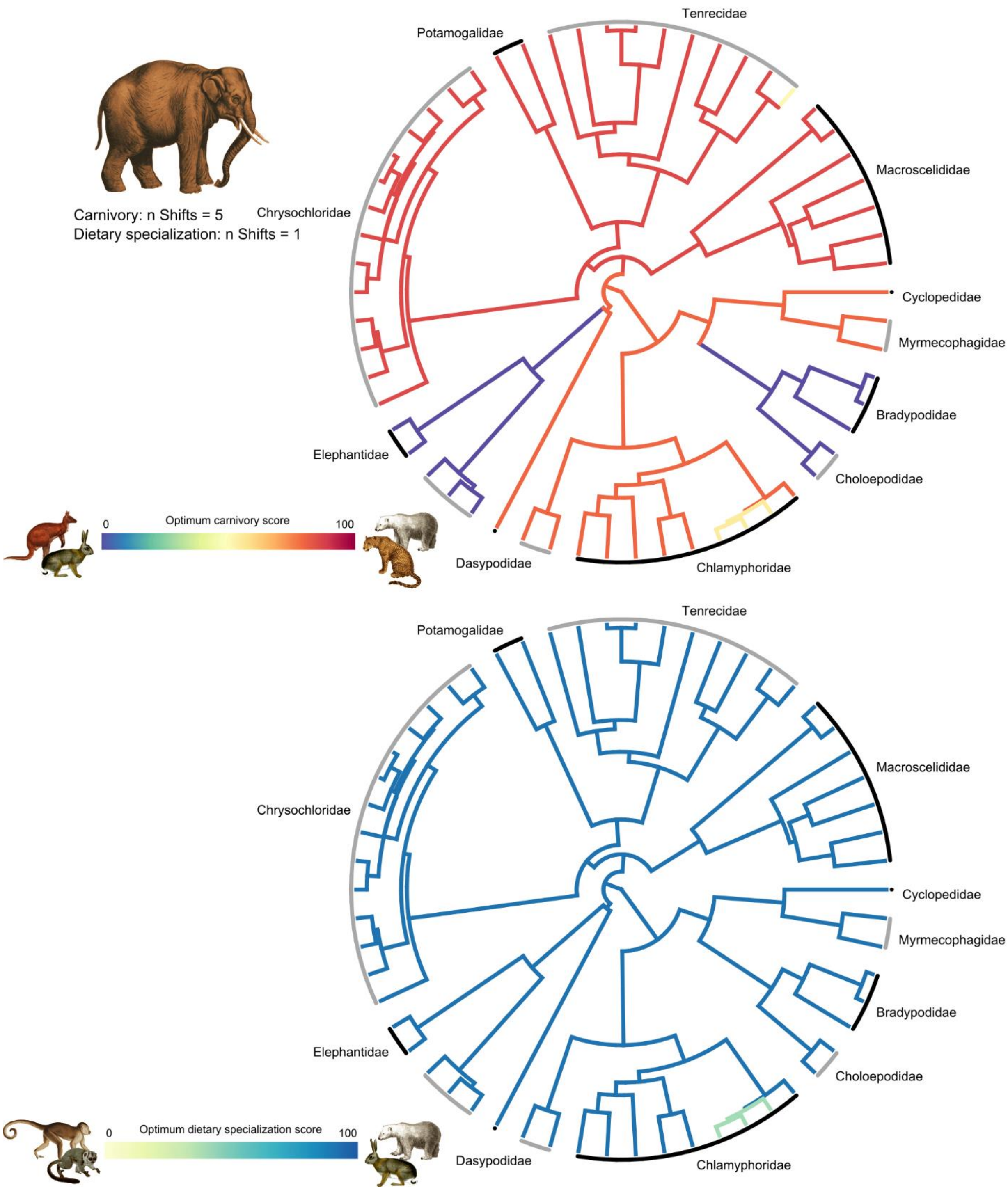

**Figure S17.** Regime shifts in optimum carnivory score (top) and optimum dietary specialization score (bottom) in Atlantogenata. Branch color indicates the value of the optimum diet score. Position of color changes represents the location of regime shifts. Images source: Wikimedia Commons.

#### Dietary regimes in Marsupialia

The carnivory regime occupied at the origin of Marsupialia is the same regime occupied at the root of Mammalia (Fig. S18). This original hypercarnivorous regime is still occupied by contemporary members of Caenolestidae, Microbiotheriidae, Notoryctidae, Thylacomyidae, and Peramelinae. At the origin of Didelphidae, there was a shift to an optimum carnivory score indicative of a diet comprising plant matter, vertebrates, and invertebrates, but with a bias towards invertivory. Subsequent carnivory regime shifts in Didelphidae include shifts to increased carnivory in the branch leading *Lestodelphis* and *Thylamys* and the branch leading to Didelphini and *Metachirus*, shifts to omnivory at the origin of Thylamyini and the branch leading to Didelphis imperfecta, and a shift to herbivory at the origin of *Caluromys*. At the origin of Dasyuromorphia, there was a shift a more carnivorous regime associated with complete carnivory and consumption of both invertebrates and vertebrates. This shift was followed by subsequent shifts to mesocarnivorous optimum diets in *Pseudantechinus bilarni* and *Parantechinus apicalis*. At the origin of Diprotodontia, there was a shift to complete herbivory. This was followed by shifts to mesoherbivory at the origins of Burramyidae and Acrobatidae, a shift to omnivory in Petauridae, and shifts to mesocarnivory in *Hypsiprymnodon* and *Acrobates*.

Fewer shifts in optimum degree of dietary specialization occurred in Marsupialia than optimum degree of carnivory (Fig. S18). The origin of Marsupialia is marked by a shift from the highly specialized regime occupied at the root of Mammalia, to a more generalized optimum diet comprising both animal and plant matter, but with a strong bias to one over the other. This regime is still occupied by Caenolestidae and Didelphidae. The origin of Australidelphia coincided with a shift to an optimum diet consistent with specialization on either plant or animal matter, and two food types within the broader category. Shifts back to regimes similar to that occupied at the origin of Marsupialia occurred at the origins of Petauridae and *Phalanger*. There was a shift to an optimal diet consisting of a single food type in *Phalanger gymnotis*. A shift to complete dietary generalism occurred at the origin of Burramyidae.

##### *Discussion*

Previous ancestral reconstruction of diet in Marsupialia have reported an insectivorous ancestor and unbroken maintenance of insectivory through to contemporary members of Australidelphia (Amador and Giannini 2021). This is reinforced with the findings of our study, as the optimum carnivory score inferred at the origin of Marsupialia could be consistent with a highly invertivorous diet, and the shift to an increased carnivory score in Dasyuromorphia is still consistent with high invertivory, although vertivory increased in prevalence.

A significant relationship between degree of carnivory and speciation rate was observed in Marsupialia. Hyperherbivores and hypercarnivores, and to some extent mesocarnivores, showed higher speciation rates than mesoherbivores and omnivores. Indeed, marsupial species that were scored as omnivorous using our carnivory variable are rare. Instead, carnivorous dasyurids and herbivorous diprotodontids comprise more of the marsupialian diversity. The radiation of these clades has been attributed to the aridification of Australia (Dodt et al. 2017; García-Navas et al. 2018), with later waves of diversification associated with the subsequent major expansion of grasslands (Kealy and Beck 2017; Celik et al. 2019). Thus, the diet-dependent speciation dynamic observed in Marsupialia may have been reinforced or driven by the impact that historical environmental change had on specific clades within Marsupialia. More work would need to be done to delineate the contributions of diet and environment to marsupial diversification.

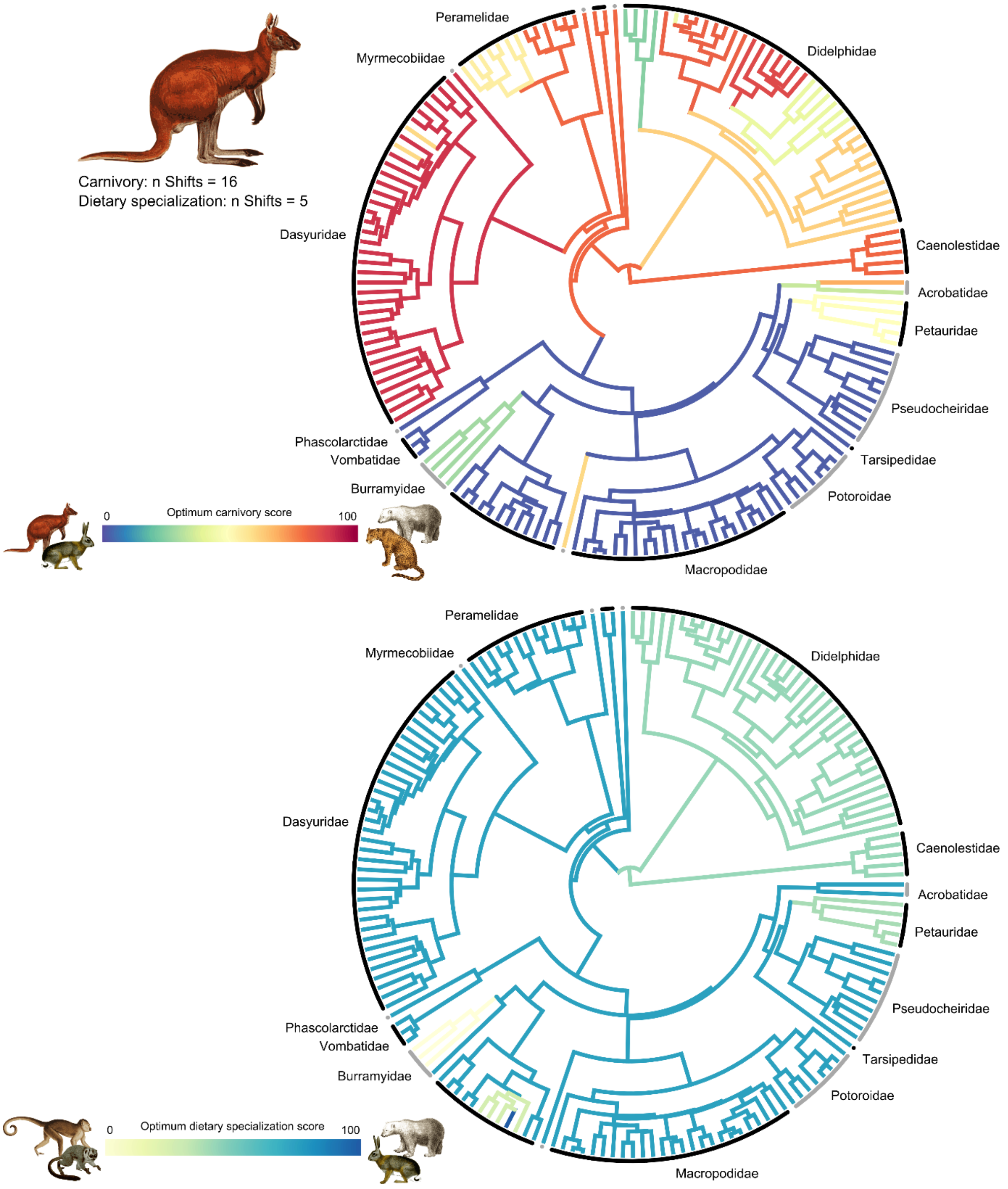

**Figure S18.** Regime shifts in optimum carnivory score (top) and optimum dietary specialization score (bottom) in Marsupialia. Branch color indicates the value of the optimum diet score. Position of color changes represents the location of regime shifts. Images source: Wikimedia Commons.

**Dietary regimes in Rodentia**

The root of Rodentia occupied the same carnivory regime that arose during the origin of Glires, with an optimum carnivory score representing complete herbivory (Fig. S19). This original regime is still occupied by some members of all rodent suborders except for Sciuromorpha. At the branch leading to Sciuromorpha, a shift towards less extreme hyperherbivory occurred. This shift in Sciuromorpha was followed by shifts towards mesoherbivory in *Heliosciurus*, *Xerus*, and Glaucomyina, shifts to omnivory in *Graphiurus* and *Glirulus*, and a single shift to mesocarnivory in *Rhinosciurus*. Shifts within Sciuromorpha back to a completely herbivorous optimum diet occurred in *Glaucomys*, *Petinomys*, and in the branch leading to *Microsciurus*, *Rheithrosciurus*, and *Sciurus*. There were no shifts away from the original rodent carnivory regime in the suborder Castorimorpha, and only a single shift in Hystricomorpha, towards a mesoherbivorous optimum diet in the branch leading to *Hoplomys*.

Most shifts in optimum carnivory score occurred in the suborder Myomorpha. In Dipodidae, there were shifts towards omnivory in *Salpingotus* and the branch leading to *Allactaga* and *Pygeretmus*. Within the family Cricetidae, shifts away from the original rodent carnivory regime occurred within Neotominae and the branch leading to Sigmodontinae. Following the shift to an omnivorous optimum diet in Neotominae, there were shifts to carnivorous optimums in the branches leading to *Onychomys* and *Scotinomys* and shifts to herbivorous optimums in the branches leading to *Baiomys* and *Megadontomys*. Following a shift towards a mesoherbivorous optimum at the origin of Sigmodontinae, there were subsequent shifts towards omnivory in the branches leading to *Sigmodon*, *Scolomys*, and the common ancestor of *Auliscomys* and *Loxodontomys*. Shifts towards varying degrees of increased carnivory occurred in *Nectomys*, *Neomicroxus*, Ichthyomyini, Akodontini, and Abrotrichini. There were shifts back to hyperherbivorous optimum diets in *Oecomys*, *Bibimys*, *Juliomys*, and at the origin of Phyllotini. Within the family Nesomyidae, there was a shift away from the original hyperherbivorous regime to a mesoherbivorous optimum diet in the branch leading to all nesomyid subfamilies, excluding Nesomyinae. This was followed by a shift towards omnivory in *Dendromus* and shifts to hyperherbivory in *Beamys* and *Delanymys*. In the family Muridae, there were shifts away from the ancestral hyperherbivorous regime to mesoherbivirous optimums at the origin of Rattini, on the branch leading to Apodemini, Murini, and Praomyini, on the branch leading to *Lorentzimys*, on the branch leading to *Stochomys* and *Hybomys*, and on the branch leading to *Apomys*, *Archboldomys*, *Chrotomys*, and *Rhynchomys*. Subsequent shifts to omnivorous optimum diets occurred in the branches leading to *Bunomys* and *Leopoldamys* and within Praomyini. Shifts to mesocarnivorous optimum diets occurred in *Melasmothrix*, *Mus triton*, the branch leading to *Praomys daltoni* and *P. derooi*, the branch leading to *Colomys*, *Myomyscus*, and *Zelotomys*. There were two shifts towards hypercarnivorous regimes within Hydromyini. Shifts back to herbivorous regimes occurred in the branches leading to *Hylomyscus* and *Chrotomys*.

Fewer shifts in degree of dietary specialization occurred in Rodentia compared to degree of carnivory (Fig. S19). At the origin of the order, there was a shift away from the highly specialized regime occupied by most of Mammalia, towards a slightly less specialized, omnivorous optimum. This specialization regime is still occupied by extant members of Anomaluromorpha, Castorimorpha, and Hystricomorpha, although shifts to increased dietary generalism occurred in the branches leading to *Anomalurus* and *Dipodomys*. At the origin of Sciuromorpha, there was a shift to an optimum specialization score consistent with an omnivorous diet comprising three different food types. Within Sciuromorpha, there were two shifts to complete dietary specialization on a single food type: at the origins of *Eupetaurus* and *Cynomys*. At the origin of Myomorpha, there was a shift to an optimum dietary specialization score consistent with a non-omnivorous diet comprising three food types, with a bias toward one foodstuff. Shifts to more generalized optimum diets occurred at the origin of Oryzomyini, the branch leading to *Rhipidomys*, the origin of Nesomyidae (exlucing Nesomyinae), the origin of Rattini, the branch leading to *Pogonomys loriae*, the branch leading to *Myodes rutilus* and *M. rufocanus*, and on the branch leading to Apodemini, Murini, and Praomyini. Shifts back to specialized optimum diets occurred in *Oecomys*, *Delanymys*, and *Paruromys*.

*Discussion*

It has been argued that the ancestral condition of Rodentia was omnivorous, and that their success as an order can be attributed to their dietary flexibility (Landry 1970). Based on the notion of an ancestral omnivorous condition, a shift away from omnivory and towards herbivory at the origin of Hystricomorpha has been suggested (Mess et al. 2008). However, our results support an optimal herbivorous condition at the origin of Rodentia, which was a regime inherited from the origin of Glires, and subsequently occupied during the formation of most rodent suborders.

Rodents represent the most speciose mammalian group, with some of the most specialized feeding apparatus within Mammalia (Cox et al. 2012; Gómez Cano et al. 2013), suggesting that diet has been important during their evolution. Despite this, diet was not found to be significantly associated with rodent diversification in the current study, or previously (Alhajeri and Steppan 2018). It was suggested by Alhajeri and Steppan (2018) that the opportunistic feeding behavior of most rodents, even those considered predominantly specialized for a given food type, renders diet classification of rodent species inaccurate. This opportunism therefore dilutes the signal of diet-dependent diversification.

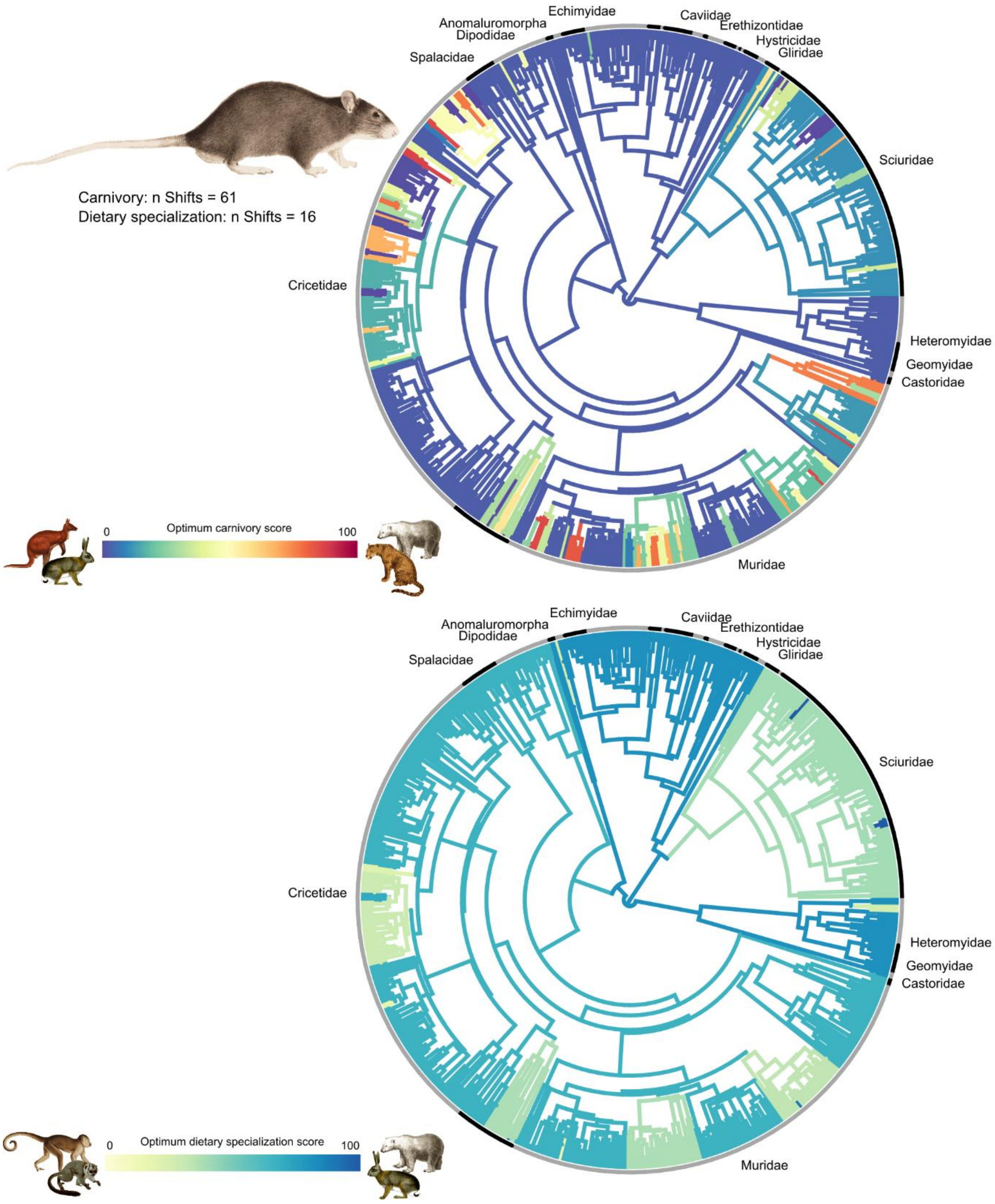

**Figure S19.** Regime shifts in optimum carnivory score (top) and optimum dietary specialization score (bottom) in Rodentia. Branch color indicates the value of the optimum diet score. Position of color changes represents the location of regime shifts. Images source: Wikimedia Commons.
